# A Suite of Foundation Models Captures the Contextual Interplay Between Codons

**DOI:** 10.1101/2024.10.10.617568

**Authors:** Mohsen Naghipourfar, Siyu Chen, Mathew K. Howard, Christian B. Macdonald, Ali Saberi, Timo Hagen, Mohammad R. K. Mofrad, Willow Coyote-Maestas, Hani Goodarzi

## Abstract

In the canonical genetic code, many amino acids are assigned more than one codon. Work by us and others has shown that the choice of these synonymous codon is not random, and carries regulatory and functional consequences. Existing protein foundation models ignore this context-dependent role of coding sequence in shaping the protein landscape of the cell. To address this gap, we introduce cdsFM, a suite of codon-resolution large language models, including both EnCodon and DeCodon models, with up to 1B parameters. Pre-trained on 60 million protein-coding sequences from more than 5,000 species, our models effectively learn the relationship between codons and amino acids, recapitualing the overall structure of the genetic code. In addition to outperforming state-of-the-art genomic foundation models in a variety of zero-shot and few-shot learning tasks, the larger pre-trained models were superior in predicting the choice of synonymous codons. To systematically assess the impact of synonymous codon choices on protein expression and our models’ ability to capture these effects, we generated a large dataset measuring overall and surface expression levels of three proteins as a function of changes in their synonymous codons. We showed that our EnCodon models could be readily fine-tuned to predict the contextual consequences of synonymous codon choices. Armed with this knowledge, we applied EnCodon to existing clinical datasets of synonymous variants, and we identified a large number of synonymous codons that are likely pathogenic, several of which we experimentally confirmed in a cellbased model. Together, our findings establish the cdsFM suite as a powerful tool for decoding the complex functional grammar underlying the choice of synonymous codons.

## 1 Introduction

The canonical genetic code, the blueprint that links nucleic acid instructions to proteins, is highly degenerate. 18 of the 20 amino acids are encoded by more than one codon. These synonymous codons were long thought to be interchangeable as they encode the same amino acid. However, codon usage bias (CUB), which refers to the non-uniform use of synonymous codons, has long been known and studied [9, 35, 43, 49, 50, 64, 73]. Increasing evidence consistently shows organism-specific [35, 43, 50] or tissue-specific [6, 20, 65] patterns of codon usage in numerous species. In addition, a number of studies revealed an important role for codon usage in the regulation of gene expression and protein folding through various mechanisms[3, 7, 8, 13, 71, 78]. To name a few, synonymous codons have been shown to be influenced by the levels of cognate tRNA and tRNA gene copy numbers and therefore differentially impact translation elongation speed [43, 50, 64]. Moreover, more than 60 synonymous variants (mutations that do not alter the amino acid sequence) associated with diseases [19, 45], and over 450 linked to tumors [72], have been reported in ClinVar, highlighting the rich information encoded within nucleotide sequences beyond what is reflected in the amino acid sequence.

Protein language models have revolutionized our understanding of protein structures and functions by learning from a large number of protein sequences [1, 30, 33, 41, 46, 56, 68]. These models have become invaluable tools in fields such as protein engineering, synthetic biology and cancer therapeutics. However, the choice of synonymous codons that encode the protein sequence falls in the blind spot of these models. By ignoring these, protein language models miss critical regulatory and structural information present in coding sequences. There has also been a growing interest in developing language models that operate directly on DNA and RNA sequences [4, 17, 29, 39, 44, 60, 62, 90]. Both genomic and protein language models leverage self-supervised learning objectives to capture the underlying biological grammar of nucleotide and protein sequences, respectively, encompassing both coding information and regulatory elements. By doing so, they can be effectively deployed in downstream tasks where labeled data are limited, such as variant effect prediction [14, 22, 38, 88], gene expression prediction [2, 61], open reading frame (ORF) localization [57], protein function prediction [47, 83], and many other applications [36, 37, 48, 79–81].

To address the gap in our understanding of codon usage and better measure the impact of synonymous codons on protein expression and function, we generated a dataset of synonymous variants across three human surface proteins, and observed that while the amino acid sequence is conserved, the choice of synonymous codons can impact overall and surface expression of proteins. This is consistent with our earlier work demonstrating the link between codon usage, tRNA abundance and translation [26]. Therefore, synonymous codons are not always interchangeable; however, the underlying contextual grammar that governs the choice synonymous codons is not well-explored. In this study, we asked whether foundation models trained on coding sequences can capture the contextual interplay of synonymous codons that gives rise to these biological variations. For this, we developed a large suite of transformer models, including encoder and decoder architectures called EnCodon and DeCodon, respectively, at various scales. We pre-trained all models on a large corpus of 60 million coding sequences (CDS) from more than 5000 species aggregated from the NCBI Genomes database (referred to as NCBI CDS throughout the paper). We studied these pre-trained models to explore the biological concepts they capture at different scales. We also evaluated their performance on a variety of zero-shot and few-shot learning tasks related to gene function and post-transcriptional regulation. We then turned our focus to tasks related to the choice of synonymous codons. Our results both reflect the importance of synonymous codons on protein expression and the ability of our pre-trained large language models to capture their influence. By applying EnCodon and DeCodon models, zero-shot, to somatic variations observed in human cancers, we have nominated many synonymous variants that are likely cancer drivers; a form of pathogenicity that current protein-based therapeutic strategies cannot address. Taken together, our findings showcase the power of EnCodon and DeCodon models to capture the context-dependent function of synonymous codons and better represent the coding sequence beyond what is captured by existing DNA and protein models.

## 2 Results

### 2.1 Leveraging self-supervised learning to pre-train EnCodon and DeCodon

Both encoder and decoder transformer architectures can be effectively applied to coding sequences. Although bidirectional encoder transformers (BERT) are frequently the preferred choice for biological sequences [18, 29, 39, 44, 62, 90], causal language models have also demonstrated remarkable zero-shot and few-shot capabilities in this field [59, 60, 70]. Given this, along with the fact that translation is inherently directional, we chose to evaluate both architectures. We developed EnCodon and DeCodon models, codon-level language models using encoder and decoder architectures, respectively. These models were pre-trained on an aggregated dataset of 60 million coding sequences from 5,000 species, primarily composed of bacterial sequences (98.7%, or 59.4 million sequences), but also including mammals and primates (Figure 1a,b,c). Based on the length distribution of the coding sequences (Figure 1c), we set the maximum context size to 2048 codons for both EnCodon and DeCodon, ensuring that over 99.5% of all sequences would fit within this context window.

**Figure 1:**
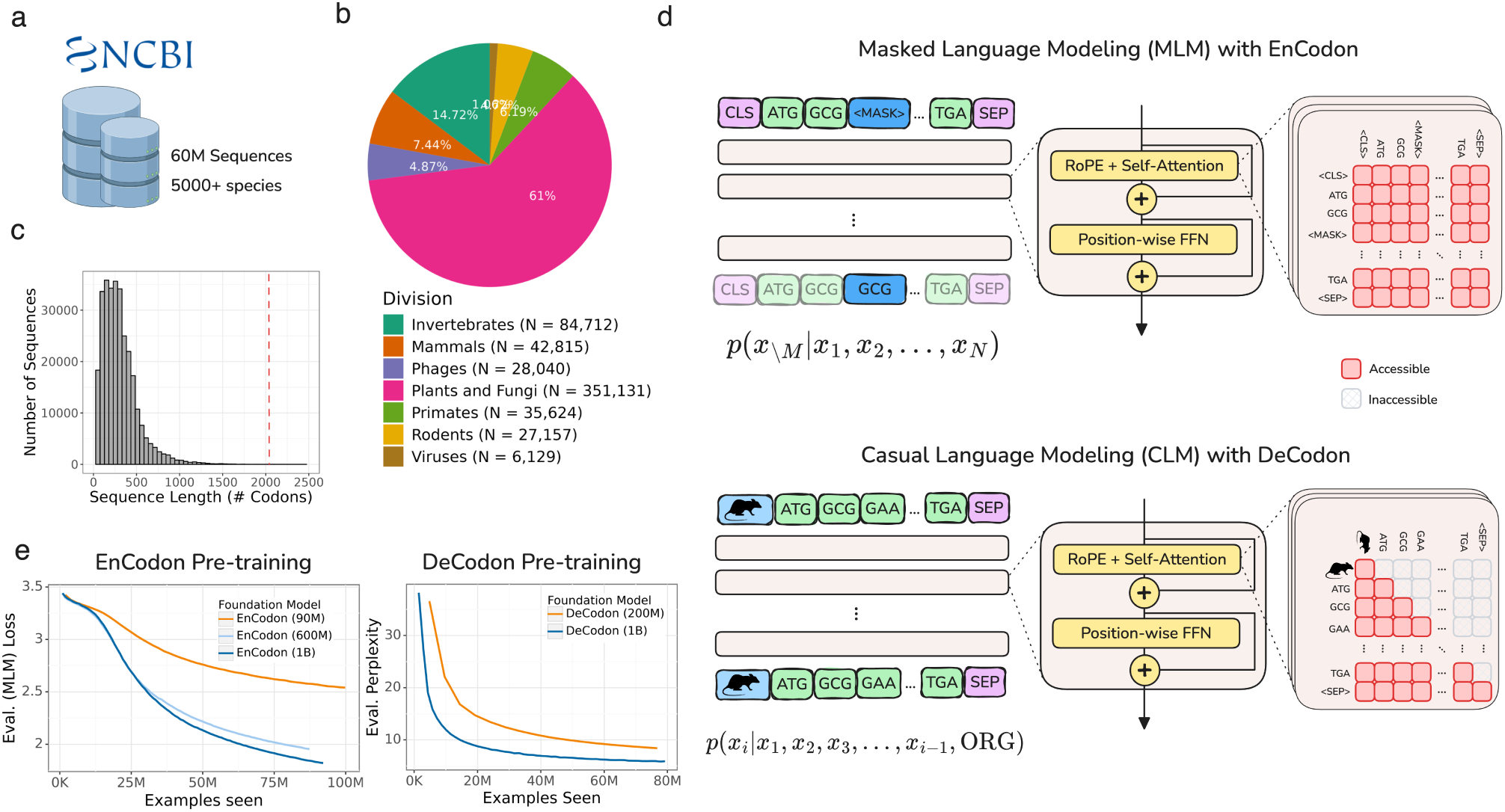
Overview of EnCodon and DeCodon: **a)** Over 60 million coding sequences from 5000 species has been extracted from NCBI Genomes database and used to pre-train EnCodon and DeCodon foundation models. **b)** An overwhelming majority of the data (98.7%) is comprised of bacterial coding sequences. Pie chart depicting division makeup of non-bacterial coding sequences in NCBI is shown. **c)** Histogram of coding sequence lengths (number of codons) in NCBI Genomes database. We used 2048 as maximum sequence length supported by EnCodon and DeCodon based taking the shown distrbution into account to cover more than 99.8% of sequences. **d)** We pretrained EnCodon using masked language modeling (MLM) objective where parts of sequences were corrupted/masked and the model has to predict the true token at the positions given the rest of tokens (i.e. context). DeCodon is a conditional generative transformer model which provides controllable coding sequence generation by querying sequence organism as the very first input token. We pre-trained DeCodon with causal (autoregressive) language modeling objective on aggregated corpus of coding sequences where each sequence is prepended with a special organism token. Rotary Positional Self-Attention was used in both EnCodon and DeCodon blocks. **e)** 3 EnCodons and 2 DeCodons, differing in scale (i.e. number of trainable parameters) have been pre-trained for more than 1,000,000 optimization steps on the aggregated corpus from NCBI Genomes database.

During pre-training, two special tokens were added to each sequence: a <CLS> token (for EnCodon) or an organism-specific token (for DeCodon) was prepended and a <SEP> token was appended (Figure 1d). EnCodon was trained using the self-supervised objective of Masked Language Modeling (MLM), where a subset of codons was randomly masked, and the model was tasked with predicting the original codons using the contextual information provided by the unmasked tokens. In contrast, DeCodon used Causal Language Modeling (CLM), where the model generated the next codon based on the preceding context (Figure 1d). To evaluate the impact of model size on performance, we pre-trained three versions of EnCodon with 80 million, 620 million, and 1 billion parameters, and two versions of DeCodon with 200 million and 1 billion parameters. As shown in Figure 1e (and Supplementary Figure 3), the larger models achieved better performance, as indicated by lower MLM loss for EnCodon and lower perplexity for DeCodon. Furthermore, we observed that the sequencing embedding space learned by all models separate by domains of life, as visualized in Supplementary Figure 1. To address the significant imbalance in the training data—where 98.7% of the data consisted of bacterial sequences, while eukaryotic sequences were underrepresented, we conducted a second stage of pre-training focused on eukaryotic sequences to better adapt the models to these coding sequences. This adaptation significantly improved performance and generalizability for eukaryotic organisms, as demonstrated in subsequent experiments (Supplementary Figure 2,4). The eukaryotic-adapted versions of each pre-trained model are denoted with a superscript “Ada”, e.g., EnCodon (1B)*^Ada^* (see Methods 5.2).

### 2.2 Codon embeddings learned by EnCodon and DeCodon reflect the structure of the canonical genetic code

To better understand the biological principles learned by EnCodon and DeCodon models, we focused on their learned codon embeddings. We assessed the relationship between codons and the amino acids they encode (Figure 2a and Supplementary Figure 5a). As shown in Figure 2b and Supplementary Figure 5b, synonymous codons are embedded substantially closer to each other relative to non-synonymous pairs. In other words, the EnCodon and DeCodon models have learned the structure of the genetic code. Beyond synonymous codon embeddings, we observed that all pre-trained EnCodon and DeCodon models exhibited a consistently strong correlation (See Figure 2c and Supplementary Figure 5c) between the pairwise cosine distance of codon embeddings and the pairwise Hamming distance of their nucleotide sequences. This notable correlation indicates that all of our models have inherently captured “load minimization” [23, 25, 58], a key property of the canonical genetic code, without explicit supervision. Load minimization refers to the ability of the canonical genetic code to reduce the detrimental effects of mutations by ensuring that common mutations are less likely to cause significant changes in protein structure or function. This is measured and demonstrated using the physicochemical properties of amino acids, such as hydrophobicity, or directly from “Point Accepted Mutation” (PAM) matrices. As shown in Figure 2d-e and Supplementary Figure 5d-e, the learned codon embeddings and the cosine distances between them very well capture this load minimization principle.

**Figure 2:**
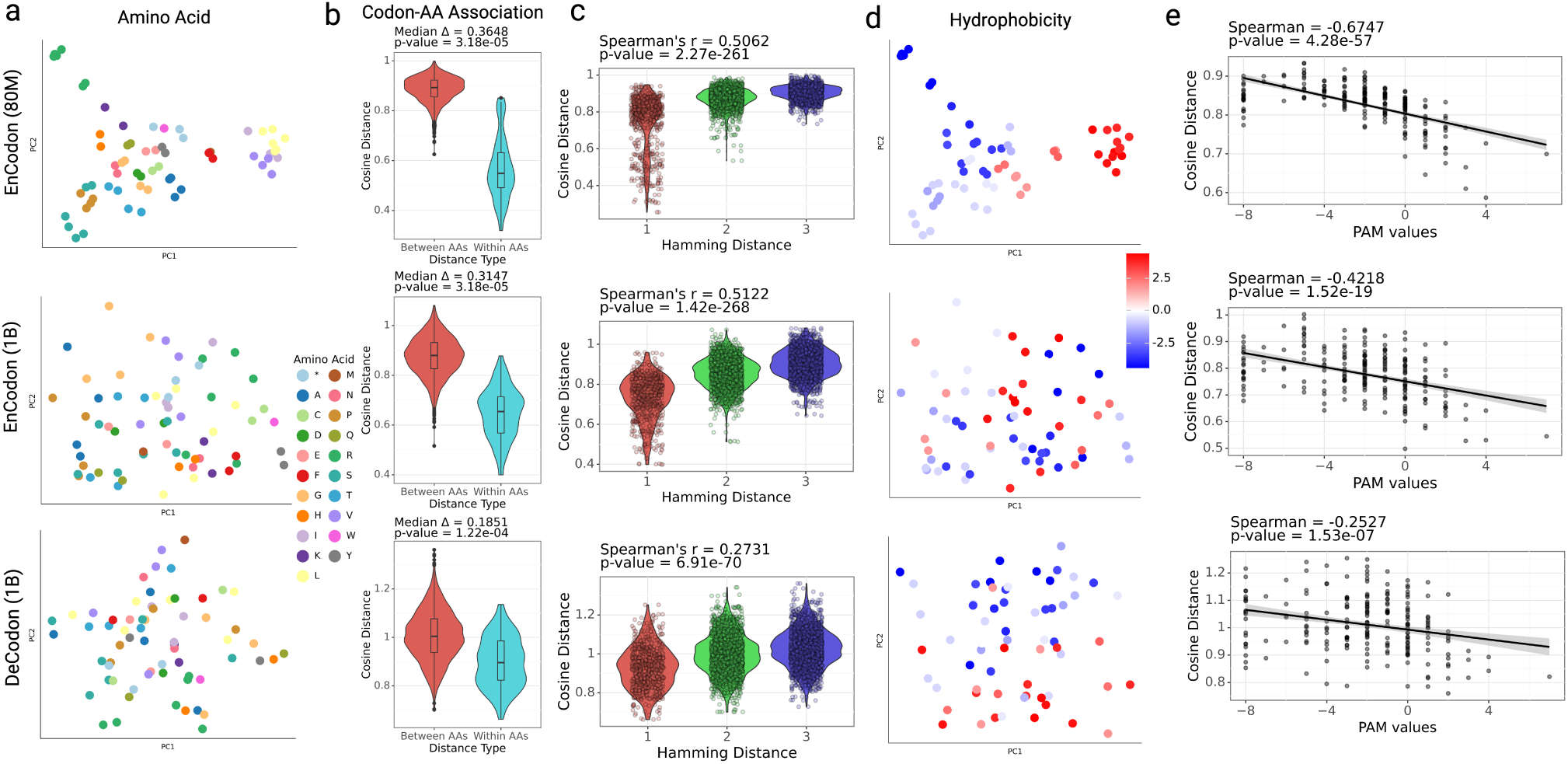
Codon Embedding Space Analysis for pre-trained EnCodons and DeCodons: **a)** PCA visualization of codon embeddings learned by EnCodon (80M), EnCodon (1B), and DeCodon (1B) colored by Amino Acid. **b)** Violin plots of two cosine distance between pairwise synonynous against non-synonymous codons for the 3 models. **c)** Violin plot of two codon distance metrics i.e. cosine distance in learned embedding space and hamming distance between codon sequences for all possible pairs of codons annotated with spearman correlation between the two metrics. **d)** PCA visualization codon embeddings colored by their corresponding amino acid’s Hydrophobicity Index. **e)** Scatter-plot of pair-wise cosine distance between amino acids and their corresponding PAM250 entry score for pretrained models.

As mentioned earlier, the larger models are more adept at distinguishing synonymous codons. This is reflected in their codon embeddings as well (See Figure 2a, Supplementary Figure 5f,g). Smaller EnCodon and DeCodon models showed better amino acid KNN purity scores (particularly for K values between 3 and 5) (Supplementary Figure 5f,g), indicating that synonymous codons are embedded closely together, and are therefore hard to distinguish. Conversely, larger models achieved significantly lower (better) Masked Language Model (MLM) losses during pre-training (See Figure 1e, Supplementary Figure 2a), by better distinguishing synonymous codons, in general moving beyond the simple structure of the genetic code. In other words, the smaller models have largely learned the mapping between the codons and the amino acids they encode, and therefore act as small protein language models with difficulty distinguishing synonymous codons. For instance, the Spearman correlation between the top 10 PC components of codon embeddings and the codon’s amino acid hydrophobicity index reveals that this information is captured in the first two PCs in smaller models. In contrast, the embedding space learned by larger models are less dominated by simple amino acid properties, with amino acid hydrophobicity captured in PC3 and beyond (See Supplementary Figure 2c). This observation highlights that larger models have learned a more sophisticated understanding of context-dependent usage of synonymous codons, and are expected to perform better in downstream tasks related to synonymous variants.

### 2.3 DeCodon generates functional organism-specific coding sequences

Since DeCodon models are generative, we sought to assess their learned knowledge of coding sequences by comparing model-generated sequences to natural ones. For this, we generated 10,000 coding sequences each for human and <u>E. coli</u>, respectively. Comparing average codon frequencies of wild-type and generated coding sequences, both DeCodon models showed strong Spearman correlations between the reference and generated sequences for both human (Figure 3a) and <u>E. coli</u> (Supplementary Figure 6c,d); two species with significantly different codon usage patterns. We then used our pre-trained EnCodon models to compare reference and generated sequence embeddings and we observed that sequences generated by DeCodon (1B) are notably better mixed with the reference cluster relative to randomly generated sequences (Figure 3b and Supplementary Figure 6a). Finally, we used a protein functional annotation tool called InterProScan to predict functional regions of the generated coding sequences. Using the computed sequence embeddings by EnCodon, we clustered the generated sequences, extracted enriched biological pathways in each cluster and annotated each cluster based on the most specific enriched pathway (Figure 3c, Supplementary Figure 6e,f,g). This observation highlights the ability of DeCodon models to generate coding sequences that capture a variety of learned protein sequence patterns.

**Figure 3:**
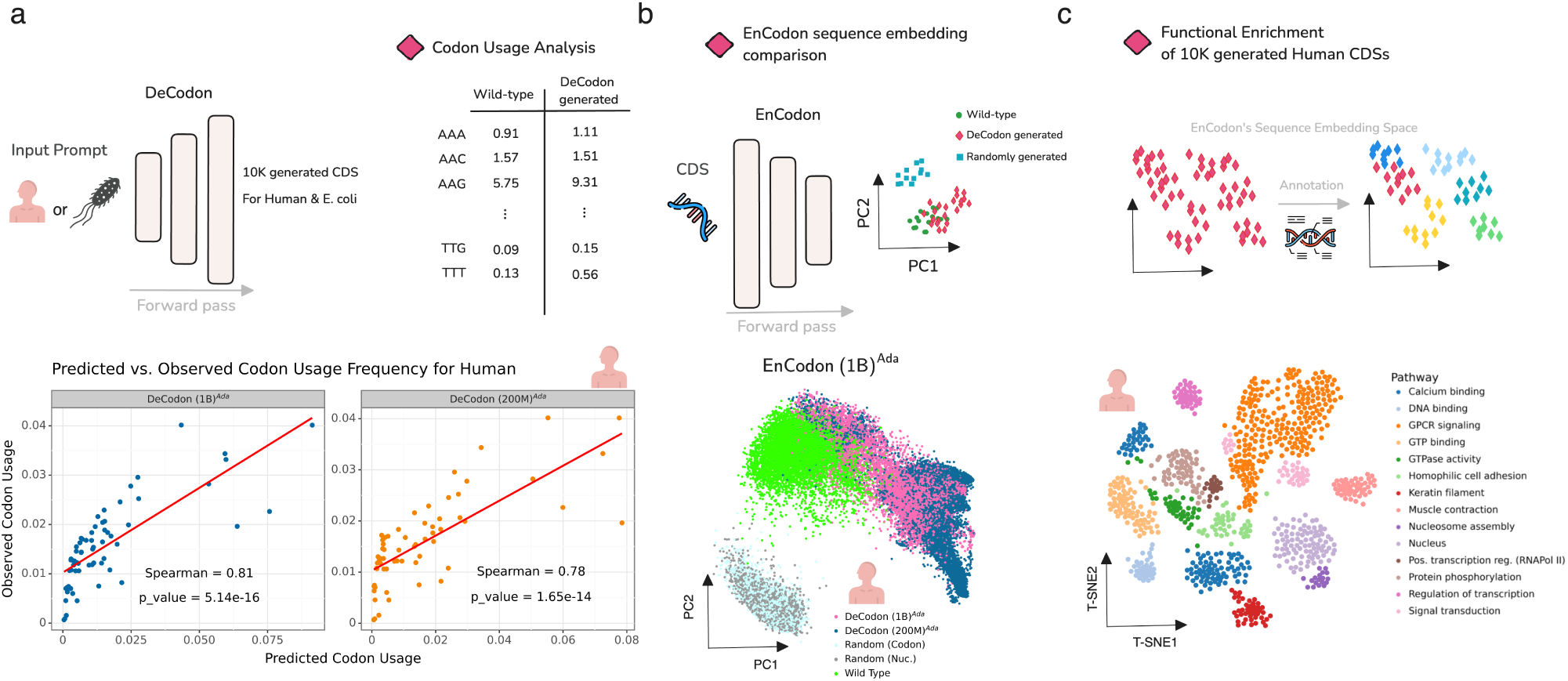
DeCodon generates functional and organism-specific coding sequences: **a)** DeCodon takes organism as input and generates a coding sequence specific to the queried species. We generated 10,000 coding sequences (CDS) for Human and E. coli species. Scatter plots of codon usage frequencies of wild-type (y-axis) and generated (x-axis) is shown for human annotated with spearman correlation and associated p-value. **b)** To further compare the generated CDS population with the wild-type, we generated two groups of randomly sampled CDSs and computed sequence embeddings of wild-type, DeCodon generated, and randomly generated groups. PCA visualization of sequence embeddings is shown for human-related coding sequences. **c)** Finally, we used protein functional annotation tools to test the functional enrichment of the sequence clusters in EnCodon embedding space. We used InterProScan to predict functional domains of human-generated CDSs by DeCodon (1B)*^Ada^*. T-SNE visualization of functionally annotated generated sequences by DeCodon (1B)*^Ada^* is shown where generated sequences were colored by their enriched biological pathway.

### 2.4 Codon Foundation Models perform zero-shot function prediction across DNA and RNA modalities

#### 2.4.1 Clinical variant effect classification in human genome

One of the key benefits of foundation models is their ability to apply their learned principles to new tasks without the need for retraining. We therefore evaluated the zero-shot capabilities of our codon language models in a variety of biologically relevant downstream tasks. As the first task, we examined the ability of our models to identify pathogenic variants annotated for the human proteome. The assessment of variant pathogenicity is considered a suitable challenge due to the large combinatorial space of all potential variants and their interactions. Furthermore, the majority of clinically analyzed variants are classified as benign, thereby limiting the number of positive instances to learn from. Recently, language models trained on corpora of nucleotide or protein sequences (exclusively utilizing self-supervised learning objectives) have demonstrated exceptional predictive performance in downstream tasks such as variant pathogenicity classification.

To perform zero-shot variant pathogenicity prediction, we calculated the pseudolikelihood (or sequence likelihood for decoder models) (See Methods 5.15.5) of 48,283 single nucleotide variants (SNVs) documented in ClinVar (see Figure 4a and Supplementary Figure 8, 9) using our language models and state-of-the-art nucleotide language models, including CaLM [62], CodonBERT [44], HyenaDNA [60], Nucleotide Transformer [18], and DNABERT 2 [90]. Upon evaluating the zero-shot pathogenicity prediction capabilities of the proposed language models on these clinically validated variants, we observed that, as expected, the Eukaryotic-adapted EnCodon and DeCodon exhibited superior performance relative to all other evaluated models (Figure 4a).

**Figure 4:**
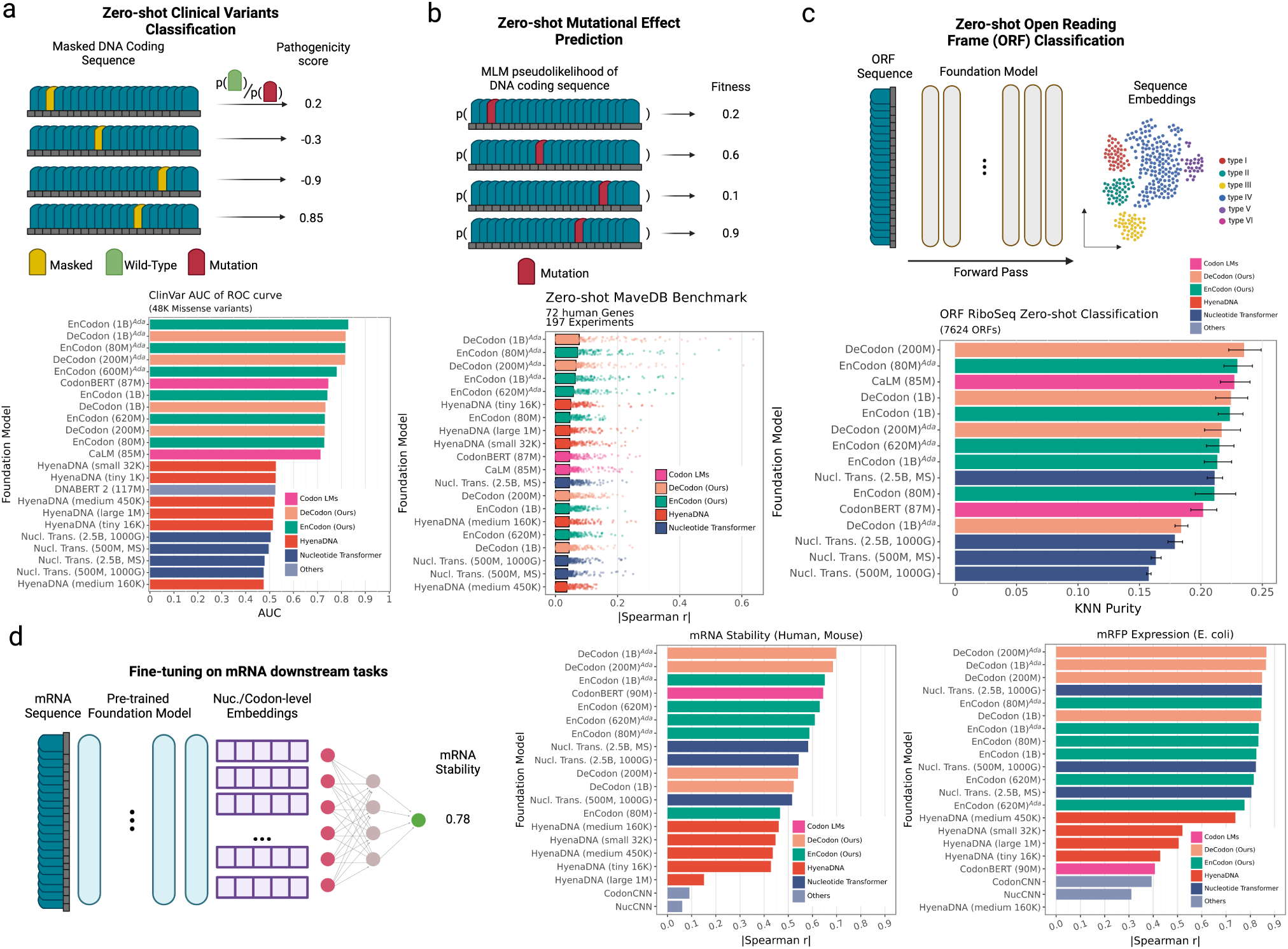
Downstream Benchmark of state-of-the-art nucleotide language models: **a)** We tested zero-shot predictive capability of the language models on 48,042 filtered variants from ClinVar. For Encoder models like EnCodon, we masked the position of variant in coding sequences and computed the log-likelihood ratio (LLR) between mutated and wild-type codon/nucleotide/token. Concerning generative models like DeCodon, we defined the pathogenicity score of variant as difference between wild-type and mutated sequence likelihoods. The Area under the ROC curve (AUC) is shown as bars for each model colored by the architecture family used in the language model. **b)** We further compared the zero-shot capability of the language models on deep mutational scan studies (DMS). Spearman correlation is reported for each model as the correlation between the reported “fitness” score and predicted sequence likelihood (pseudolikelihood for encoder models). 197 Human DMS studies across 72 genes were used to assess the language models shown in the barplot where each dot is a single human DMS study. The bars are colored according to the architecture family used in the language model. **c)** We further tested the foundation models’ discriminative capability of sequence embedding space in localization of open-reading-frames (ORFs). We use a recently published collection of 7624 open-reading frames (ORFs) where sequences were annotated based on their genomic location. KNN Purity scores (using different Ks) of the extracted sequence embeddings were shown in bar plot where bars represent means showing as bars and lines as standard deviations. **d)** We used LoRA [34] technique for parameter-efficient fine-tuning of the language models on mRNA-related downstream tasks namely mRNA stability and mRFP expression prediction. For each downstream task, a barplot of Spearman correlation of the hold-out test set for all fine-tuned models on mRFP Expression (E. coli) and mRNA stability (Human and Mouse) tasks.

#### 2.4.2 Predicting mutational effects on protein function measured by deep mutational scan (DMS) studies

We applied our CDS foundation models to predict the effect of mutations on protein function, utilizing data from deep mutational scanning (DMS) studies. These studies introduce comprehensive sets of mutations to protein sequences and experimentally measure their impact on fitness – a study-specific metric that quantifies protein functionality ([21]). To predict experimental fitness scores, similar to the previous task, we used pseudolikelihoods (or likelihoods for autoregressive models, See Methods 5.15.5) of codons or nucleotides generated by the models (Figure 4b). Our analysis was restricted to DMS studies that provided nucleotide information for wild-type sequences and their corresponding mutations. Therefore, we evaluated the foundation models on DMS studies across 7 different organisms (including human, <u>E. coli</u>, house mouse, chicken, etc), covering approximately 58,262 and 16,403 mutations, respectively (Figure 4b, Supplementary Figure 10 and 11).

The Eukaryotic-adapted codon language models consistently placed as the top five performers, showcasing the highest correlations for both human and <u>E. coli</u> datasets (Figure 4b, Supplementary Figure 10). This outcome indicates the efficacy of our adaptation technique and the versatility of our pre-trained models to specific organismal contexts. All coding sequence models, including CodonBERT (which ranked 6th), surpassed HyenaDNA and nucleotide transformer models. As expected, both the adapted and pretrained versions of our codon foundation models excelled in the <u>E. coli</u> benchmark (Supplementary Figure 10a). When comparing EnCodons with DeCodons, we consistently noted an increase in performance with larger DeCodon models, although the scale of improvement for both models were the same when the model size was fixed. These findings underscore the potential of our codon-level large language models in forecasting mutational impacts on protein functionality.

#### 2.4.3 Evaluation of Nucleotide Language Models for Human Open Reading Frame (ORF) Classification

We next sought to focus on tasks centered on the translational capacity of the transcriptome. Ribosome profiling (Ribo-seq) has significantly broadened our understanding of the translational output of the cell by uncovering numerous open reading frames (ORFs) in regions previously believed to be untranslated, such as long non-coding RNAs (lncRNAs) and untranslated regions (UTRs) of protein-coding genes [5, 12, 40, 52, 66, 84]. Mudge et al. [57] described a standardized, spatially-classified, and filtered collection of translated ORFs, integrating previously reported ORF databases. Given the multiple interpretative approaches possible for ORFs, the sequences were systematically classified based on their spatial relationships with existing gene annotations. Our objective was to evaluate our models’ predictive accuracy in classifying ORFs without further supervision (i.e., training). To this end, we employed the standardized catalog of 7,264 human Ribo-seq ORFs to assess the proficiency of embeddings from various models in ORF classification (Figure 4c). Accordingly, we reported the statistics of purity scores of the K-Nearest Neighbors (KNN) algorithm for each model across different K values, ranging from 5 to 50 (Figure 4d). The results indicated that our codon foundation models consistently outperformed all other tested language models, with DeCodon (200M) achieving the highest average KNN purity score of 24.57%, whereas the Nucleotide Transformer (2.5B, multi-species) achieved an average score of 21.45%, representing the best performance among the existing state-of-the-art language models.

#### 2.4.4 Codon language models show superior performance on various mRNA downstream tasks

We further compared the foundation models’ performance across a few supervised mRNA prediction tasks namely mRNA stability (Human and Mouse) [2] and mRFP expression (E. coli) [61] prediction. Similar to zero-shot benchmarks, we benchmarked Nucleotide Transformer[11], HyenaDNA [60], CaLM [62], and CodonBERT [44] as prior methods alongside with two convolutional baselines i.e. NucCNN and CodonCNN (See 5.12). We used low-rank adaptation technique (LoRA) [34] to fine-tune the language models in the downstream tasks. Our benchmark shown in Figure 4d shows our EnCodon and DeCodon models outperform their counterparts in both mRNA stability and mRFP expression prediction tasks. More specifically, DeCodon (1B)*^Ada^* showed 5% improvement in mRNA stability prediction over Nucleotide Transformer (2^nd^ best model), underscoring its superior understanding of mRNA dynamics.

### 2.5 EnCodon captures the effect of synonymous codon mutations on protein abundance levels

While most synonymous variations are considered neutral, there is mounting evidence that synonymous variations can also carry functional consequences [19, 71, 78]. Being able to discriminate between synonymous codons and better capture their possible context-dependent role in the translational output was a major motivation behind training EnCodon and DeCodon models. However, there are few instances of functional synonymous variants, and we currently lack large datasets evaluating synonymous codons. To address this gap, we generated a dataset of 1054 synonymous coding variants across three membrane proteins with directly comparable phenotypes, namely SLC22A1, GPR68, and KCNJ2 [32, 51, 89]. While all datasets are on membrane proteins, they represent distinct protein architectures and functions, including a tetrameric 2 transmembrane cardiac potassium channel associated with defects associated with developmental disorders [82]; KCNJ2, a monomeric 12 transmembrane polyspecific cation transporter that has genetics variants associated with differential effects on drug metabolism [27]; and a proton sensing receptor, GPR68, a seven transmembrane with variants associated with chemotherapy-induced neuropathy [42]. We then systematically measured the overall abundance and surface expression of these variants. To test our models’ predictive performance, we held out 513 variants from SLC22A1 from the training dataset of the foundation models. Figure 5a depicts the fine-tuning procedure where the model takes wild-type sequence as input and log-likelihood ratio between wild-type and mutated codon at the mutated position is used to predict the abundance level of the protein (See 5.14). In other words, without the need to introduce any additional parameters to the pre-trained model, we repurposed the MLM head to model the variant’s effect. Comparing different pre-trained EnCodons (with or without adaptation) on holdout test variants in SLC22A1, consistent improvement in test performance is observed as the model gets larger (Supplementary Figure 12a and Supplementary Figure 13). As expected, the eukaryotic adaptation of the EnCodons also outperformed their non-adapted same-size model resulting in adapted EnCodon (1B) being the best performer which performed well across experimented proteins and hold-out test set (Figure 5b,c, Supplementary Figure 12a,b). The fine-tuned EnCodon model performs slightly better on surface expression than protein abundance in the one gene, KCNJ2, in which we have both measures for abundance and surface expression. This is likely because surface expression is dependent upon protein abundance but also includes effects that change whether the protein makes it to the surface.

**Figure 5:**
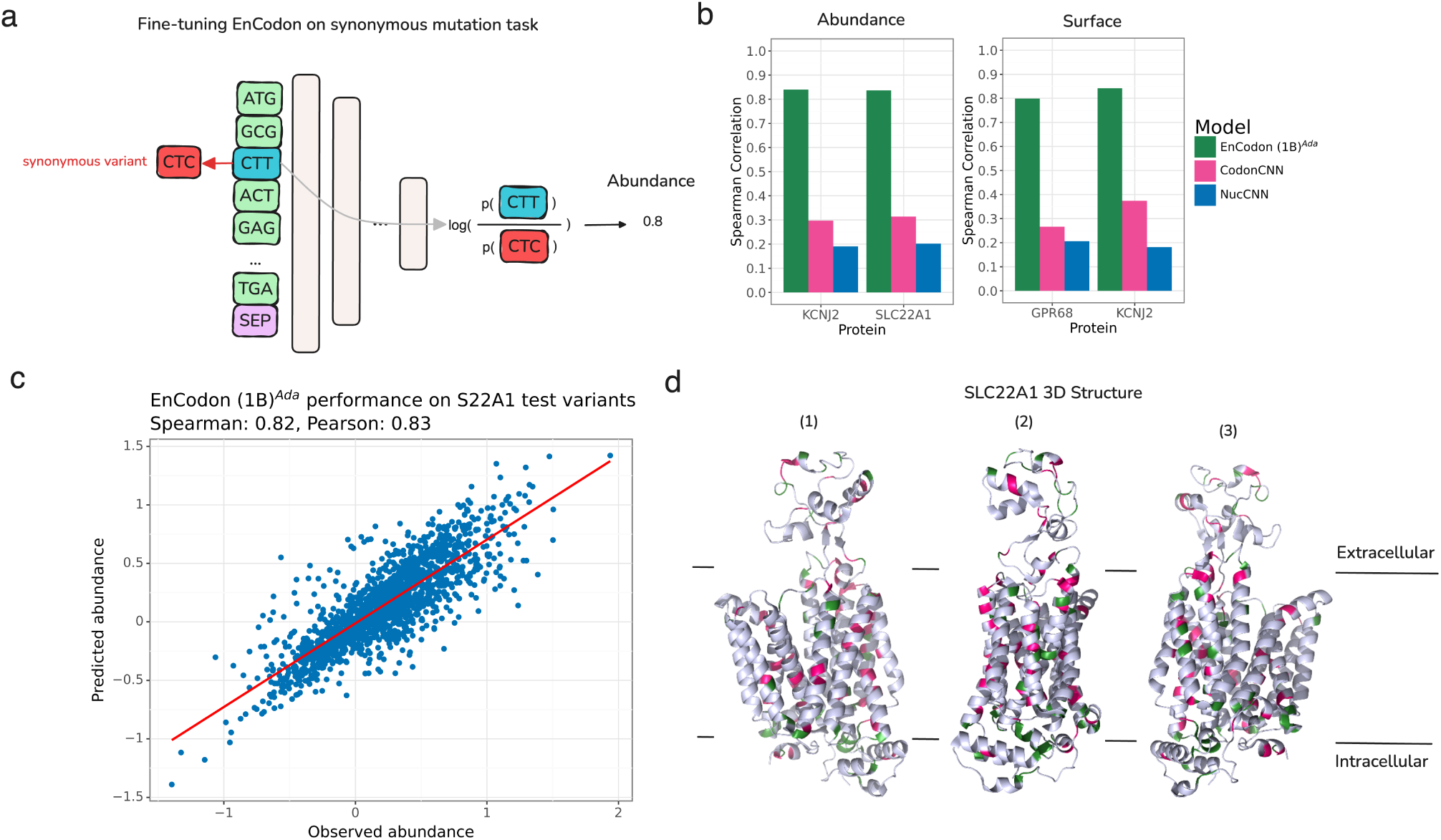
EnCodon (1B)*^Ada^* generalizes well across unseen synonymous variants in membrane proteins. **a)** We re-purposed the pre-trained language modeling classifier head for synonymous mutation effect modeling. Specifically, given a synonymous codon variant, we first compute the codon likelihoods (i.e. logits) for the variant’s position. Next, the log-ratio of wild-type codon against mutated codon is considered as the final variant’s effect prediction – protein abundance level or surface expression in this experiment. Notably, using no additional weights, the mutation’s effect on protein’s abundance measurement is modeled as the log-likelihood ratio between mutated and wild-type codon given the *wild-type* coding sequence in input. **b)** Spearman correlation between predicted and observed abundance (left bar plot) or surface expression (right bar plot) were shown for test synonymous variants in KCNJ2, SLC22A1, and GPR68 proteins. **c)** An test set of synonymous variants applied on SLC22A1 were held-out from the training data of the EnCodons. The scatter plot of predicted vs. observed abundance is shown for eukaryotic adapted EnCodon (1B) which showed as the top-performed compared to other fine-tuned EnCodons. **d)** After fine-tuning, we performed in-silico synonymous mutagenesis with the best-performing EnCodon model. We selected “critical” synonymous variants for which the predicted abundance was above 95-th (green) or below the 5-th quantile (pink). Next, extracted SLC22A1 extreme variants were overlayed in the protein’s 3D structure which is shown from 3 different angles.

We employed our top-performing fine-tuned EnCodon model to conduct in-silico synonymous codon mutagenesis, predicting the abundance levels for all possible synonymous coding variants within the protein sequences. Variants with exceptionally high or low abundance scores (i.e., critical variants) were selected for further analysis. To evaluate the spatial distribution of these critical variants within the 3D structures of the proteins studied, we applied Ripley’s *K* function [67], a well-known function that determines a spatial pattern (i.e. random, dispersed, or clustered) of certain points (in 3D structure) at a certain distance cut-off (See Methods 5.16). More specifically, the results revealed significant clustering of the identified “critical” variants for KCNJ2 and SLC22A1 at distances below 10 Å and 8 Å, respectively (Supplementary Figure 12d). The calculated p-values from Ripley’s test, compared against a null distribution (see Methods 5.16), indicate regions where the observed clustering significantly deviates from being randomly spread (i.e., p < 0.05, Supplementary Figure 12d).

Using a 9Å cutoff radius in the Ripley’s *K* function test, we demonstrated significant clustering of extreme variants in the 3D structures of SLC22A1 and KCNJ2 (see Methods 5.16). Despite having no information on protein folding and structure, the “critical” variants identified by EnCodon are predominantly located in functional elements of the 3D structure, such as *α*-helices or *β*-sheets, emphasizing the sequence-structure relationships that EnCodon captures in its learned representations (Figure 5d, Supplementary Figure 12c). The significant clustering observed at these short distances suggests that these variants are not randomly distributed.

### 2.6 Leveraging CDS foundation models to nominate and validate novel pathogenic synonymous variants

As previously shown and demonstrated here, synonymous codon usage can impact protein structure and gene expression through effects on translation efficiency, kinetics, elongation, mRNA stability, and co-translational protein folding [3, 7, 8, 13, 43]. Given the zero-shot performance of our CDS foundation models in predicting variant pathogenicity, and the ability of our models in distinguishing synonymous codons in mutational scans in the previous section, we set out to nominate previously unknown pathogenic synonymous variants. To this end, we computed pathogenicity scores (see Methods 5.15.5) for all synonymous coding variants in ClinVar that were previously labeled as variants of uncertain significance (VUS), using our best-performing codon foundation models (in zero-shot benchmarks), including EnCodon (1B)*^Ada^* and DeCodon (1B)*^Ada^*Figure 6a). We then selected four synonymous coding variants with extremely low predicted pathogenicity scores (less than the 5th percentile, i.e., within the highly pathogenic region) and two control variants with extremely high predicted scores (greater than the 95th percentile, i.e., within the highly benign region) for experimental testing (see Methods 5.8). Our experimental results demonstrated significant changes in protein expression levels for two of the four nominated synonymous variants with extreme predicted pathogenic scores by our DeCodon (1B)*^Ada^* model: NM_006412.4(AGPAT2):c.702C>T and NM_002872.5(RAC2):c.501C>T (Figure 6b,c and Supplementary Figure 12a). These findings provide compelling evidence not only revealing the post-translational roles of synonymous codon variants but also underscoring the broader significance of these variants in regulating protein expression. Furthermore, this demonstrates the caability of our DeCodon models to capture such nuanced patterns in synonymous codon usage, emphasizing the potential utility of our model in guiding future research and clinical interpretations of synonymous variants, offering new insights into their functional consequences.

**Figure 6:**
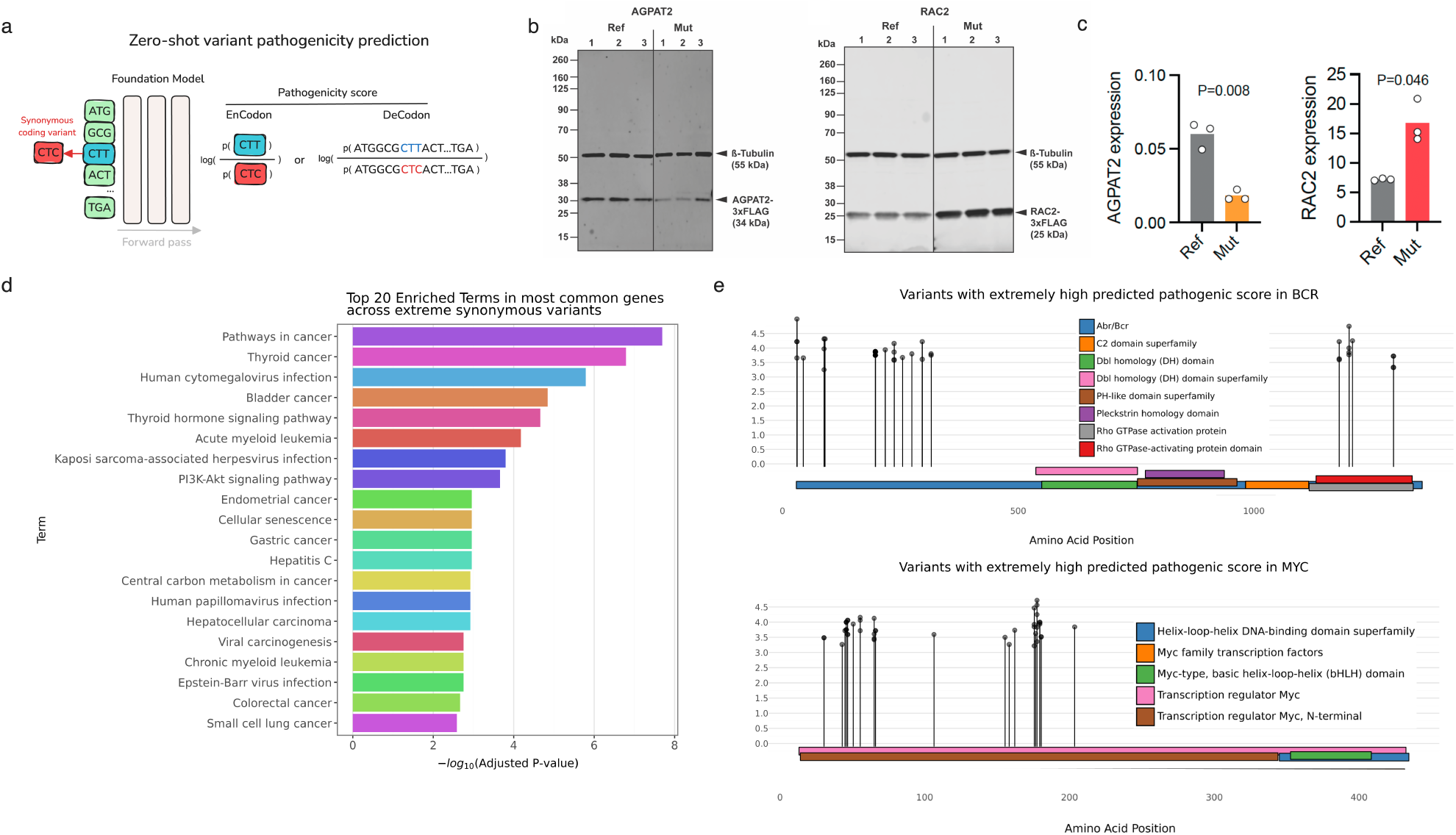
Pathogenic synonymous variants predicted by codon foundation models cause notable changes in gene expression levels: **a)** We use different scoring fashions for pathogenicity of synonymous variants. Specifically, we report log likelihood ratio between wild-type and mutated codons for EnCodon and log ratio of wild-type and mutated sequence likelihoods for DeCodon models. To nominate variants for experimental validation, We fetched all synonymous variants from ClinVar and COSMIC Census Mutations databases and computed pathogenicity scores for all of our pre-trained EnCodons and DeCodons. Next, we selected synonymous variants with extremely pathogenic scores (above 99th quantile of the distribution). **b)** Experimental measurement of gene expression levels for AGPAT2 and RAC2 are shown for wild-type vs. mutated coding sequence. Each dot denotes an individual replicates and bars show the average expression across replicates. **c)** Immunoblots of FLAG-tagged AGPAT2 and RAC2 protein variants expressed in HEK293T cells, showing three biological replicates for both wild-type (Ref) and mutated (Mut) sequences. *β*-tubulin signal is used as a loading control. **d)** Geneset Enrichment Analysis of top 20 abundant genes with highest number of “pathogenic” variants with extreme scores. **d)** Lollipop plot of extreme variants predicted for BCR and MYC are shown across protein sequence annotated by functional domains extracted from InterPro.

#### 2.6.1 Pathogenic synonymous variants identified by cdsFMs are predominantly enriched in cancer-related genes

To expand our dataset of predicted synonymous variants, we scored COSMIC Census Mutation database [75], focusing on those with extreme pathogenic variants predicted by all our CDS foundation models (See Figure 6a). Specifically, we isolated 3365 synonymous coding variants with extremely high pathogenicity scores and performed gene-set enrichment analysis on their associated genes. Given that COSMOS reports on somatic variations in cancer, we expect identifying synonymous mutations that are enriched in known drivers. As expected, the analysis revealed a significant enrichment in cancer-related pathways (Figure 6c), underscoring the potential impact of these synonymous variants on cancer biology.

For instance, BCR and MYC, which are well-established oncogenic drivers across many cancers [10, 24, 28, 31, 74], harbor 64 of these extreme variants. Lollipop plots, annotated with InterPro domains, illustrate the distribution and functional implications of these variants. The observed spatial clustering of these synonymous mutations further highlights their biological function on the expression of these proteins (Figure 6d, Supplementary Figure 14b,c,d,e,f). Using this approach, we have significantly expanded the list of likely pathogenic synonymous mutations in cancer.

## Discussion

Synonymous codons are often underappreciated and underexplored. Although the vast majority of synonymous variants are silent, many are likely pathogenic and functional. Similarly, synonymous codons are not interchangeable in synthetic biology and can be used to tune the expression of the same underlying protein. The absence of reliable machine learning models to help elucidate the context-dependent role of codons was the main motivation for building EnCodon and DeCodon models. As we embarked on this, other groups have also released codon-resolution or codon-aware models. CodonBERT, with its ability to handle codon-level inputs, has shown strong potential in certain tasks, particularly those involving sequence-level transformations. cdsBERT, on the other hand, has demonstrated proficiency in capturing protein-level semantic information from coding sequences. However, these models are an order of magnitude smaller than our largest models, are trained on fewer tokens, and are limited to the BERT architecture. As we demonstrated here, the larger models are crucial to capture the context dependence of synonymous codons. More importantly, the DeCodon models perform superior to EnCodon models in a number of key downstream tasks.

In this study, we have demonstrated the versatility and effectiveness of our suite of codon-based foundation models, EnCodon and DeCodon, in comparison to a wide range of genomic language models of various scales (i.e., differing in the number of trainable parameters) across a variety of downstream tasks relevant to coding sequences. Synonymous codons fall in the blind spot of protein language models and as we demonstrated here, genomic foundation models have not capture the codon structure in the coding regions. This is likely because coding sequences are a small minority of sequences that the genomic models are trained on. Here, we showed that codon-resolution large-scale language models extend beyond the basic codon-amino acid associations to capture more intricate codon-codon relationships, which proved highly beneficial for tasks like synonymous variant effect prediction.

In this study, we successfully confirmed two out of four synonymous variants predicted as pathogenic by our EnCodon and DeCodon models. This 50% confirmation rate is likely an underestimate given the variety of ways beyond expression by which synonymous variants may impact protein function, e.g. by impacting the folding dynamics. Our broader data generation effort for measuring the impact of synonymous codons on protein expression, further highlights the importance of capturing the functional consequences synonymous variants. Our results not only highlight the potential of EnCodon and DeCodon in practical genomic applications but also provide additional evidence for the heterogeneous effects of synonymous codons on gene regulation and expression.

In this study, we included both masked and causal language modeling, and found that the latter showed superior performance particularly in downstream few-shot tasks. This suggests that the causal modeling approach of DeCodon offers more flexible and informative context-aware representations. At first glance, this may be counterintuitive as having access to the entirety of the sequence context should provide more information. However, other models, such as Evo [59] or LoRNA*^SH^* [70], have similarly shown success in learning biological sequences via causal language modeling. The added benefit of these models is that they are generative in nature, and the coding sequences they generate can be studied to further understand the biological concepts the models have internalized.

In conclusion, our suite of codon foundation models, namely EnCodon and DeCodon, provides a powerful toolkit for advancing our understanding of synonymous codons. Given the challenges of generating deep mutational scanning at scale for synonymous codons, these models are essential for better nomination of likely functional variants. This knowledge, and the extent to which it impacts human diseases, will also reshape how we consider therapeutic modalities, as some mutations will not have an impact on protein sequence, yet it can still impact expression level or activity.

## 3 Code Availability

Code and models are accessible at https://github.com/goodarzilab/cdsFM. Additionally, pre-trained models have been made available on HuggingFace [87].

## 4 Acknowledgments

We thank Brian Plosky and Chiara Ricci-Tam for their valuable comments and insights during the preparation of this manuscript. HG is an Arc Core Investigator and this study was in part supported by the Arc Institute. WCM acknowledges support from a Howard Hughes Medical Institute Hanna Gray Fellowship, and UCSF Quantitative Biosciences Institute Fellowship, and is a Chan Zuckerberg Biohub San Francisco Investigator. MKH and CBM are supported by NIGMS 5T32GM139786 and 1F32GM152977.

---

**Supplementary Table 1.** Hyperparameters used for each of our pre-trained codon foundation models (cdsFMs). Emb. dim.: codon-level embedding dimensionality, Inter. dim.: Intermediate layers’ dimensionality, LR: learning rate, WD: weight decay

| Foundation Model | # layers | Emb. dim. | Inter. dim. | LR | WD | warmup steps |
| --- | --- | --- | --- | --- | --- | --- |
| EnCodon (80M) | 12 | 1024 | 2048 | 1e-4 | 1e-2 | 10,000 |
| EnCodon (620M) | 12 | 2048 | 8192 | 5e-5 | 1e-2 | 10,000 |
| EnCodon (1B) | 18 | 2048 | 8192 | 1e-5 | 1e-2 | 10,000 |
| DeCodon (200M) | 12 | 1024 | 2048 | 1e-4 | 1e-2 | 10,000 |
| DeCodon (1B) | 18 | 2048 | 8192 | 1e-5 | 1e-2 | 10,000 |

**Supplementary Figure 1:**
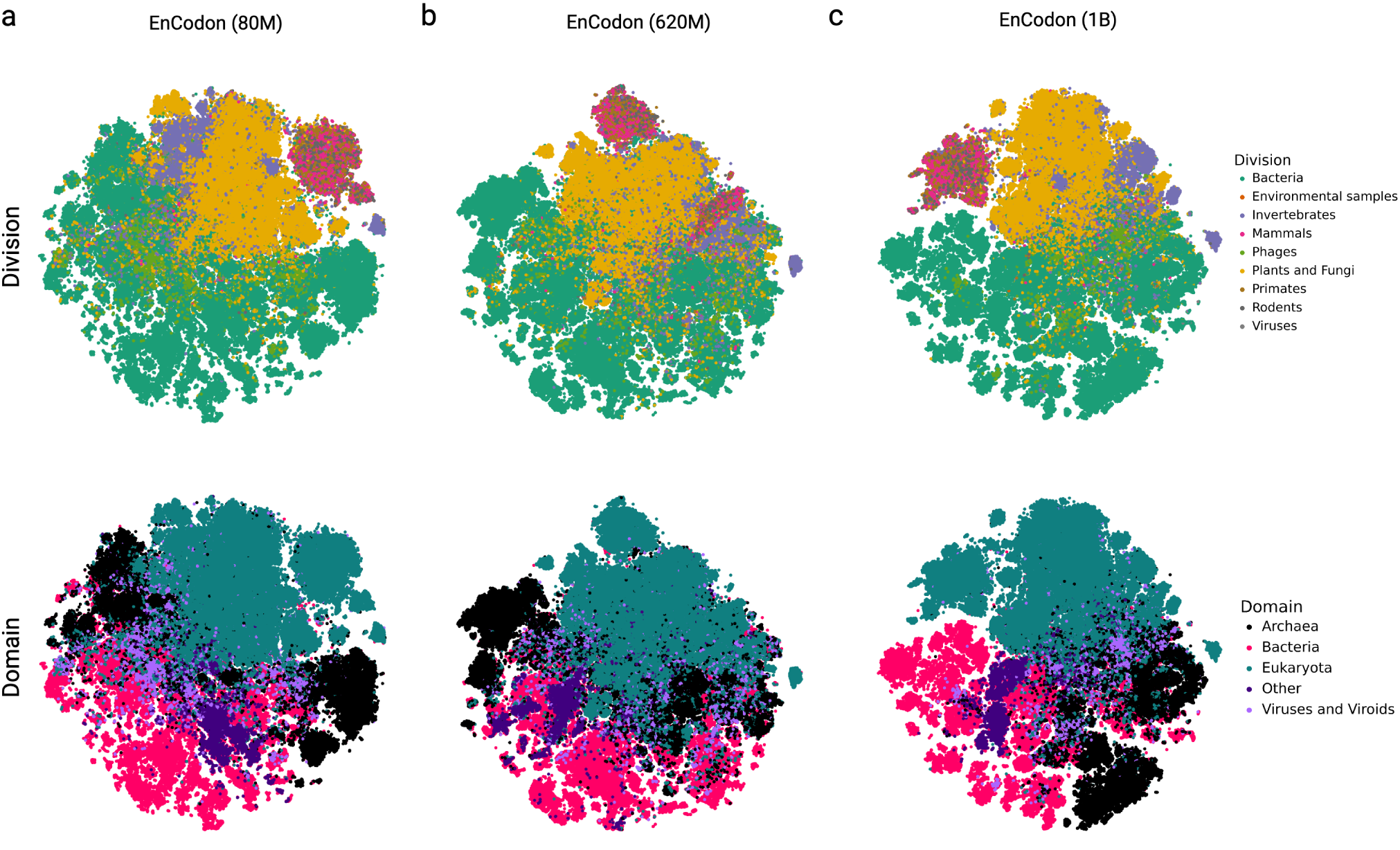
T-SNE visualization of sequence embedding space learned by a) EnCodon (80M), b) EnCodon (620M), and c) EnCodon (1B) where each dot is a sequence and they are colored by sequence’s organism division (top row) and domain (bottom row).

**Supplementary Figure 2:**
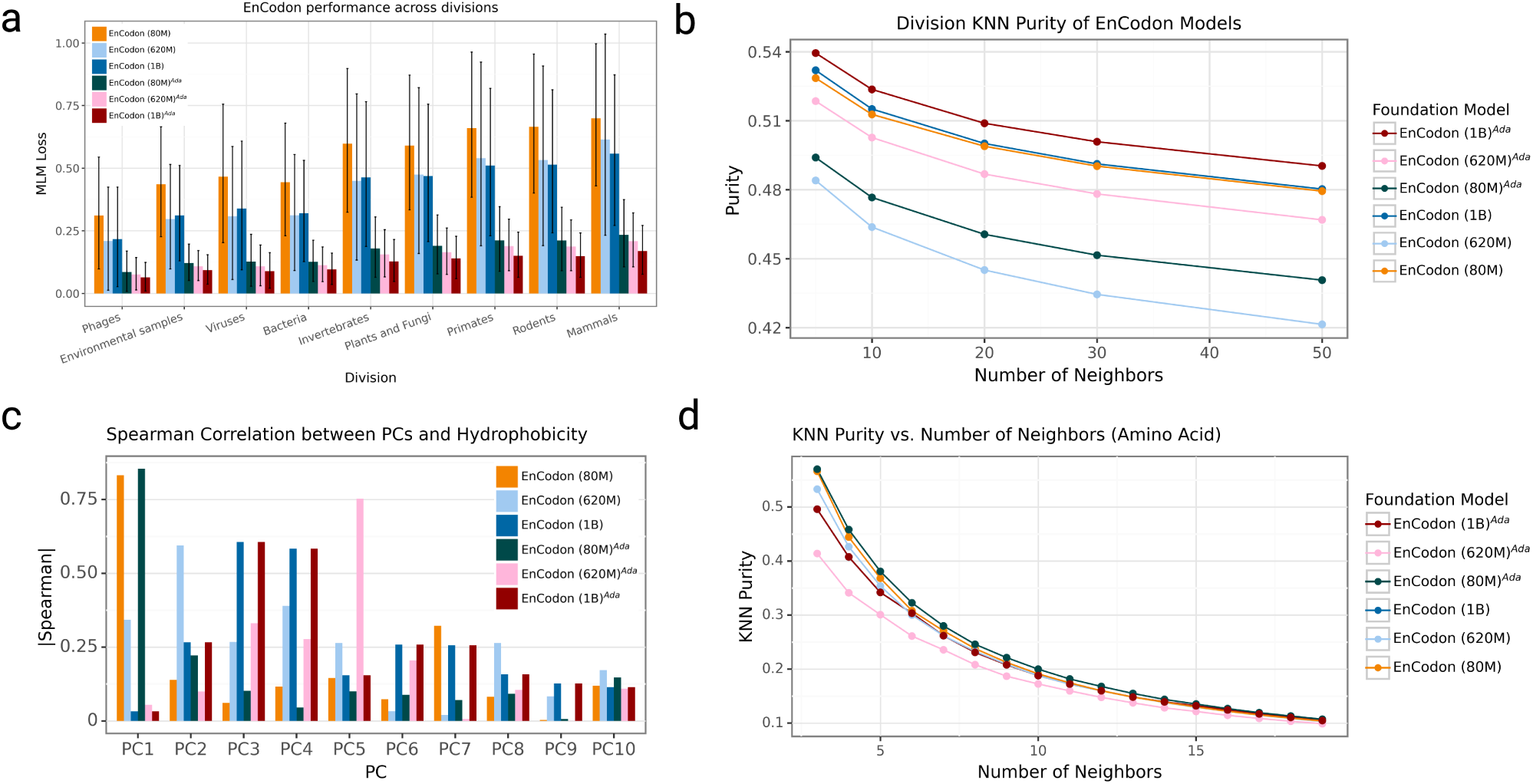
**a)** Representation of MLM loss distribution for pre-trained and adapted EnCodon models across taxonomy divisions with mean bars and standard error lines. **b)** Scatter plots of KNN Purity scores against numbers of nearest neighbors, using organisms’ Division as clustering labels. **c)** Spearman correlations bar plot between the top 10 principal components (PC) of the pretrained/adapted EnCodons and the hydrophobicity index of codon’s amino acid. **d)** KNN Purity scores of the codon embedding space of EnCodons with amino acid labels against the number of neighbors (K).

**Supplementary Figure 3:**
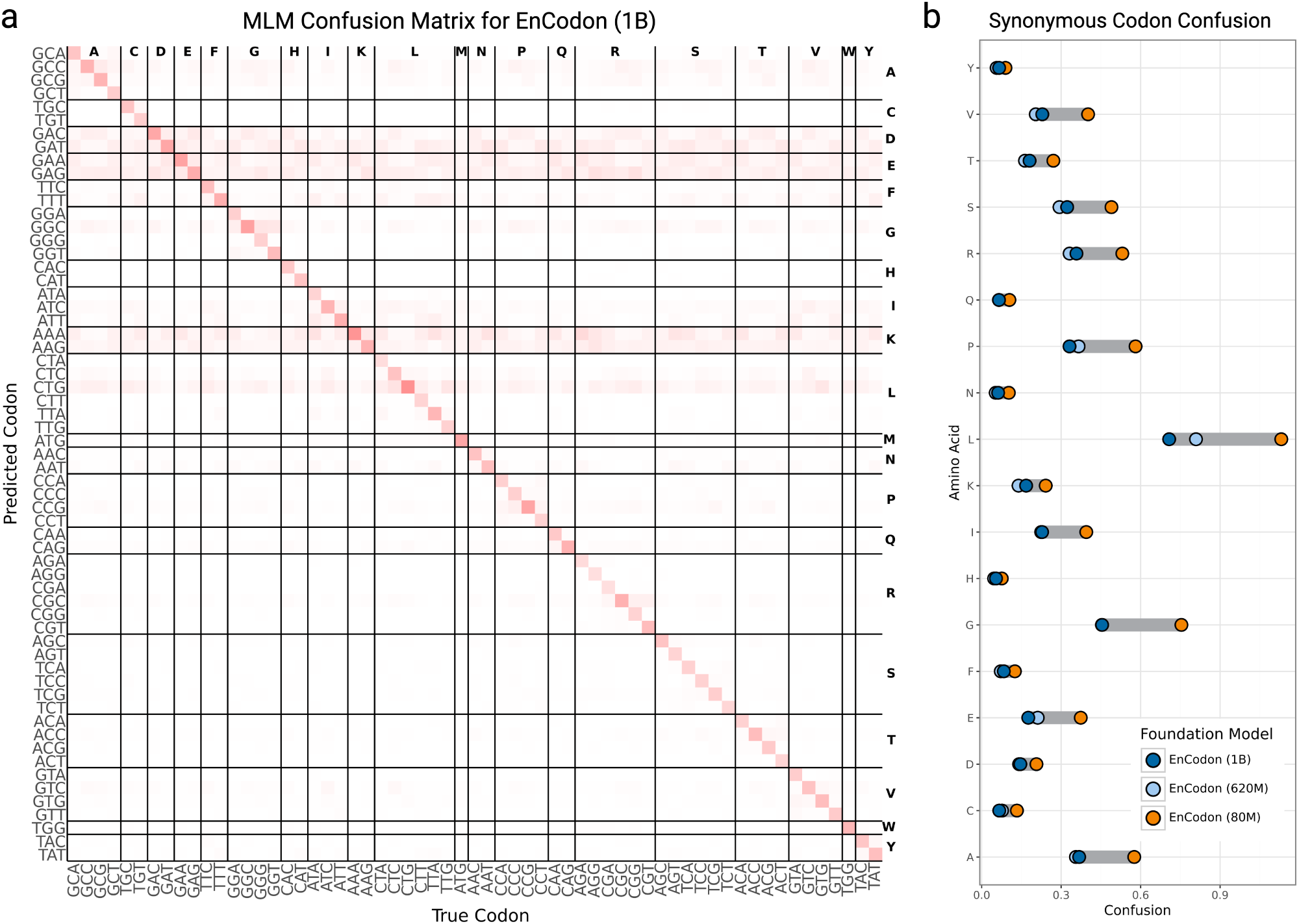
**a)** Masked language modeling confusion matrix of pre-trained EnCodon (1B) model. We use sequences in the pre-training test split and randomly masked each codon in the sequence with 0.15 probability. The shown confusion matrix is computed from EnCodon’s prediction on the masked positions. **b)** Difference plot of synonymous codon confusion per amino acid is shown for the purpose of comparing pre-trained EnCodons – EnCodon (80M), EnCodon (620M), and EnCodon(1B).

**Supplementary Figure 4:**
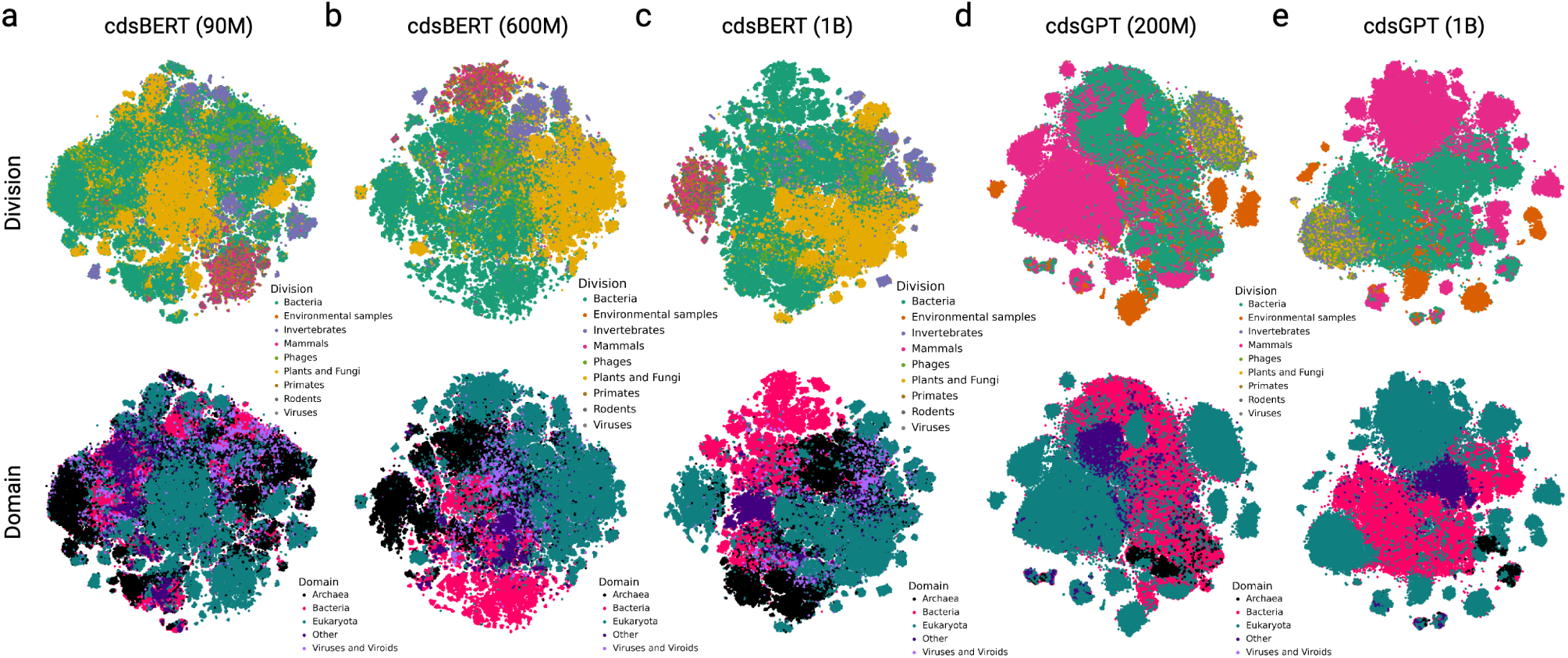
T-SNE visualization of sequence embedding space learned performing eukaryotic adaptation on **a)** EnCodon (80M), **b)** EnCodon (620M), **c)** EnCodon (1B), **d)** DeCodon (200M), and **e)** DeCodon (1B) where each dot is a sequence and they are colored by sequence’s organism division (top row) and domain (bottom row).

**Supplementary Figure 5:**
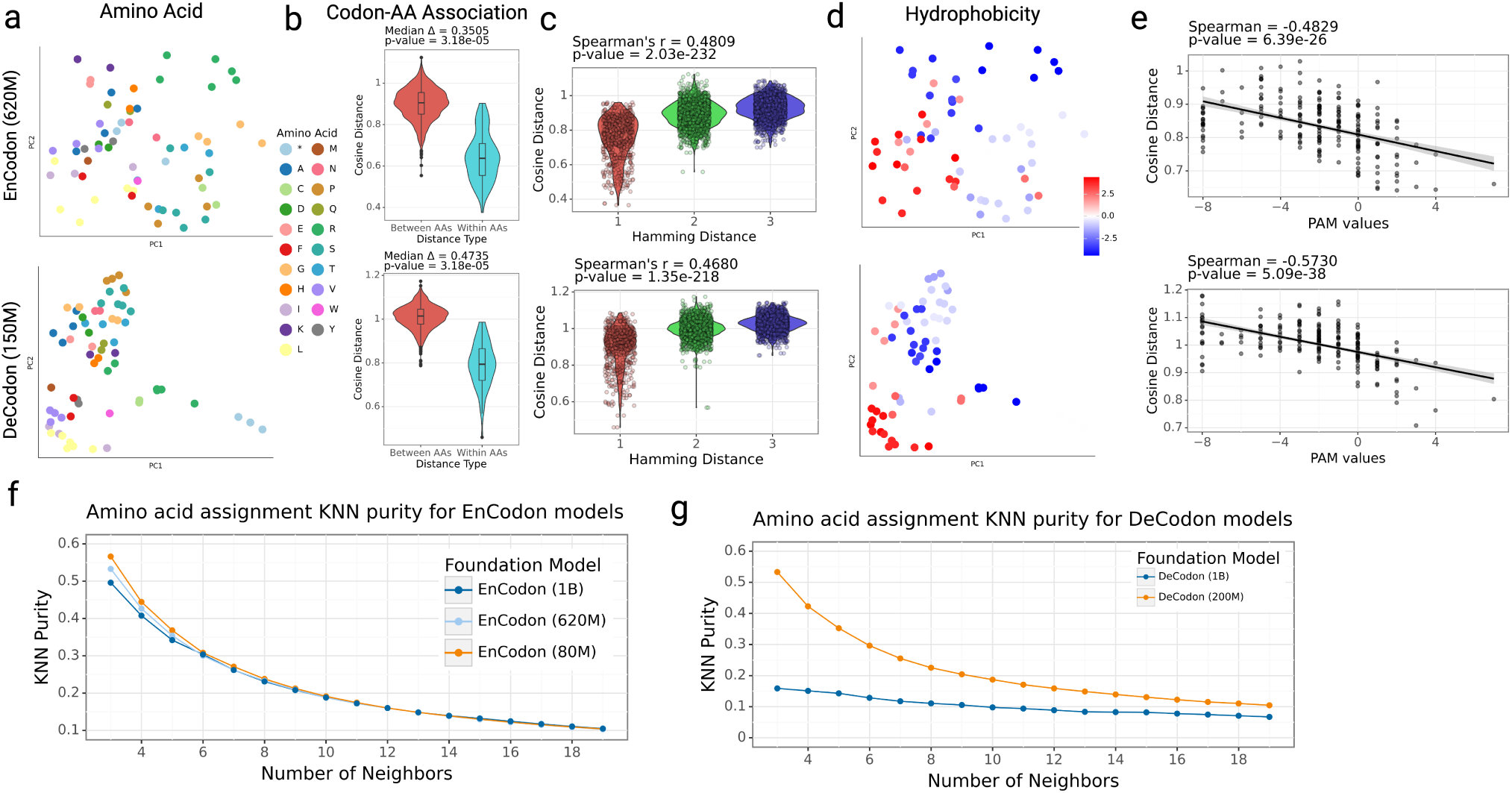
Codon Embedding Space Analysis for pre-trained EnCodons and DeCodons: **a)** PCA visualization of codon embeddings learned by EnCodon (620M) and DeCodon (150M) colored by Amino Acid. **b)** Violin plots of two cosine distance between pairwise synonynous against non-synonymous codons. **c)** Violin plot of two codon distance metrics i.e. cosine distance in learned embedding space and hamming distance between codon sequences for all possible pairs of codons annotated with spearman correlation between the two metrics. **d)** PCA visualization codon embeddings colored by amino acid’s Hydrophobicity Index. **e)** Scatter-plot of pair-wise cosine distance between amino acids and their corresponding PAM250 entry score for pre-trained models. Scatter plot of KNN purity scores of clusters of synonymous codons in learned codon embedding space by **f)** EnCodon and **g)** DeCodon models against different numbers of neighbors.

**Supplementary Figure 6:**
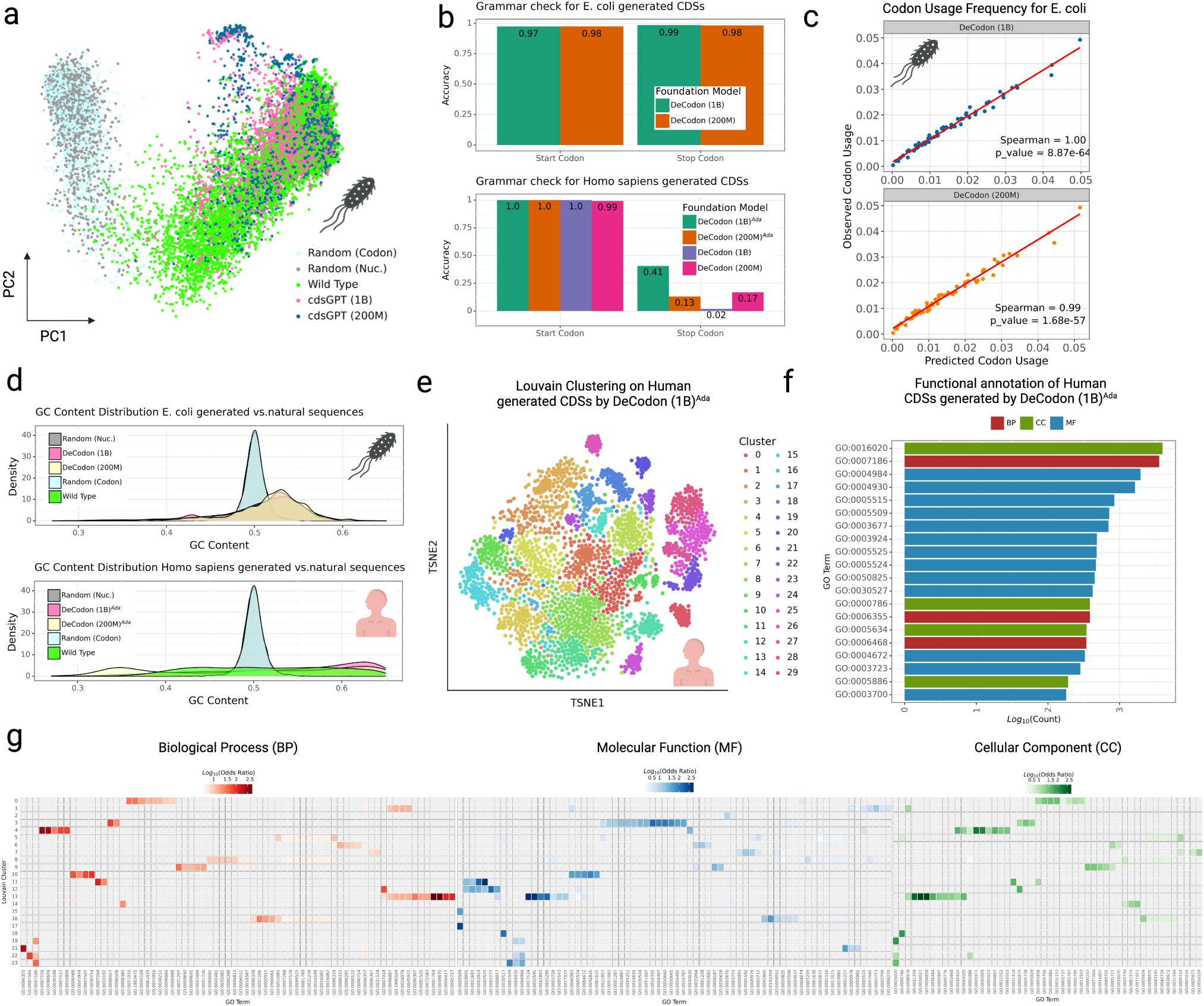
Analysis of DeCodon-generated coding sequences: **a)** PCA visualization of EnCodon (1B)*^Ada^*’s sequence embedding space to compared generated coding sequences with wild-type and random cohorts in E. coli. **b)** Accuracy bar plots of start and stop codon grammar checks on the generated coding sequences for human and E. coli. **c)** Scatter plot of observed codon usage (in wild-type sequences) against codon usage in generated coding sequences (x-axis) for E. coli. **d)** Comparison of the GC content distribution between generated sequences and natural coding sequences from E. coli (top) and human (bottom), where each distribution is compared with two sets of 10K randomly generated sequences. **e)** Louvain clustering performed on 10,000 sequences generated by DeCodon (1B)*^Ada^*, with a t-SNE visualization colored by cluster ID. **f)** Functional region prediction of generated sequences using InterPro and PANTHER, highlighting the top 20 most common Gene Ontology (GO) terms as bars representing log-transformed number of annotated sequences colored by their namespace. **g)** Fisher’s Exact Test for GO term enrichment across Louvain clusters, with a heatmap showing significant enrichments (*p*_*adjusted <* 0.05) based on the GO namespaces: biological process (BP), molecular function (MF), and cellular component (CC).

**Supplementary Figure 7:**
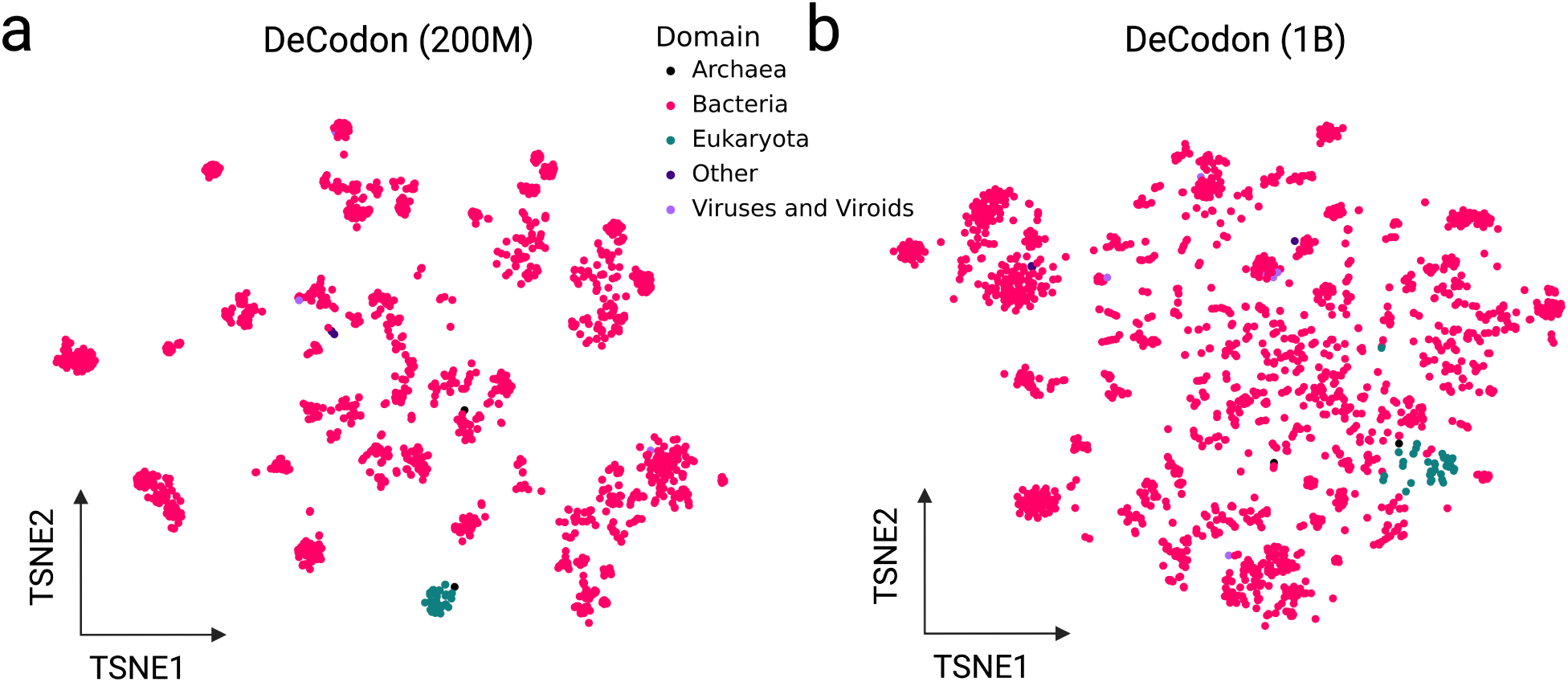
DeCodon organism embedding space: PCA visualization of pre-trained DeCodon’s organism embedding space for **a)** DeCodon (200M) and **b)** DeCodon (1B)

**Supplementary Figure 8:**
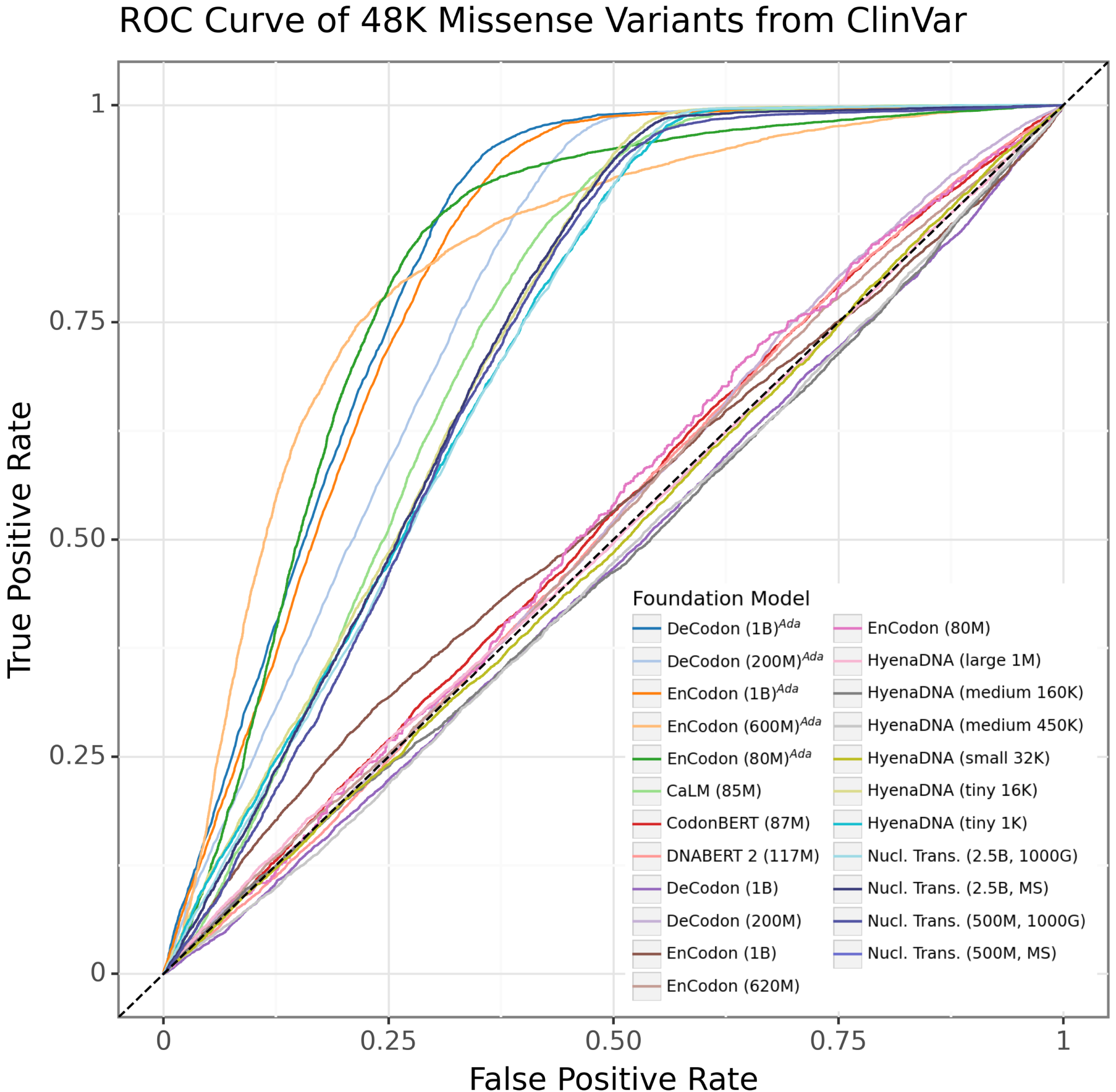
Receiver Operating Characteristic (ROC) curve depicting the True Positive Rate (TPR, y-axis) versus the False Positive Rate (FPR, x-axis) for the foundation models evaluated in predicting ClinVar variant pathogenicity. The comparison was standardized by calculating the TPR and FPR on a common set of 48,000 shared missense variants across all models.

**Supplementary Figure 9:**
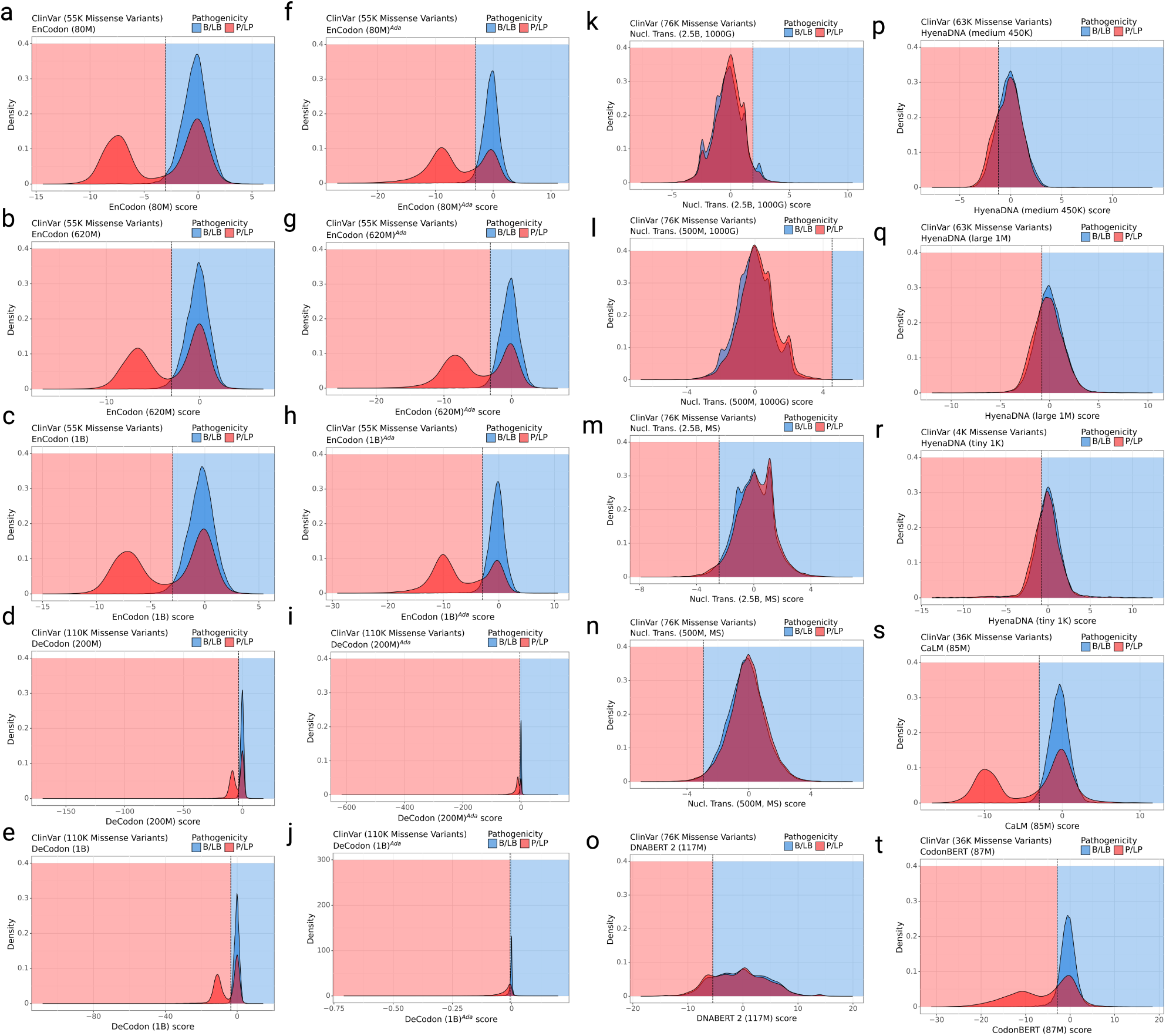
Distribution of missense variant scores for all the models used in the zero-shot ClinVar benchmark. Missense Variant scores distribution colored by the consequence of the variant where P/LP and B/LB represents Pathogenic/Likely Pathogenic and Benign/Likely Benign variants. The score distribution is shown for **a)** EnCodon (80M), **b)** EnCodon (620M), **c)** EnCodon (1B), **d)** DeCodon (200M), **e)** DeCodon (1B), **f)** EnCodon (80M)*^Ada^*, **g)** EnCodon (620M)*^Ada^*, **h)** EnCodon (1B)*^Ada^*, **i)** DeCodon (200M)*^Ada^*, **j)** DeCodon (1B)*^Ada^*, **k)** Nucleotide Transformer (2.5B, 1000G), **l)** Nucleotide Transformer (500M, 1000G), **m)** Nucleotide Transformer (2.5B, MS), **n)** Nucleotide Transformer (500M, MS), **o)** DNABERT 2 (117M), **p)** HyenaDNA (medium 450K), **q)** HyenaDNA (large 1M), **r)** HyenaDNA (tiny 1K), **s)** CaLM (85M), and **t)** CodonBERT (87M).

**Supplementary Figure 10:**
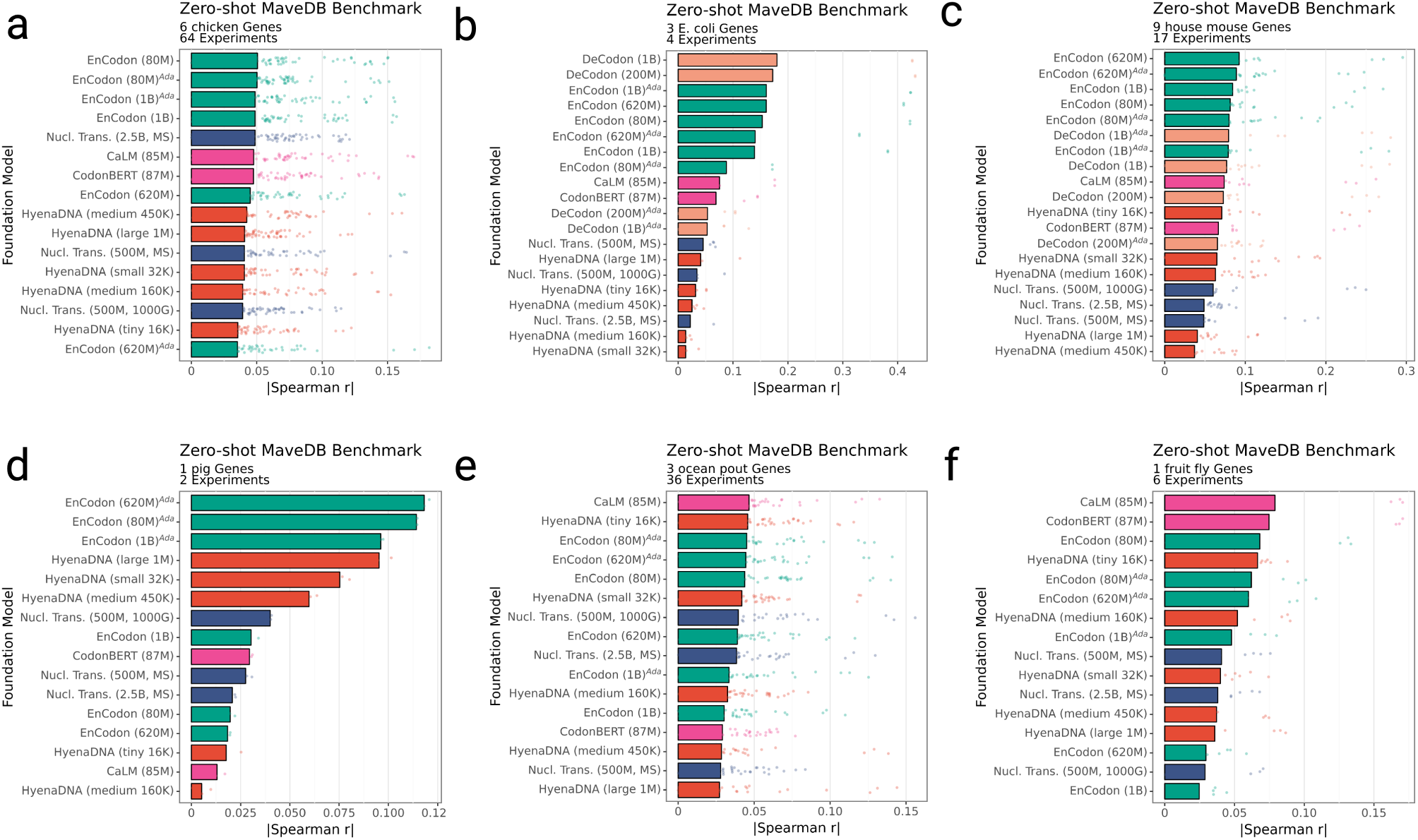
Distribution of absolute Spearman correlations (x-axis) for the tested foundation models (y-axis) across different organisms in the Zero-shot MaveDB benchmark: **a)** Chicken, **b)** E. coli, **c)** House mouse, **d)** Pig, **e)** Ocean pout, and **f)** Fruit fly.

**Supplementary Figure 11:**
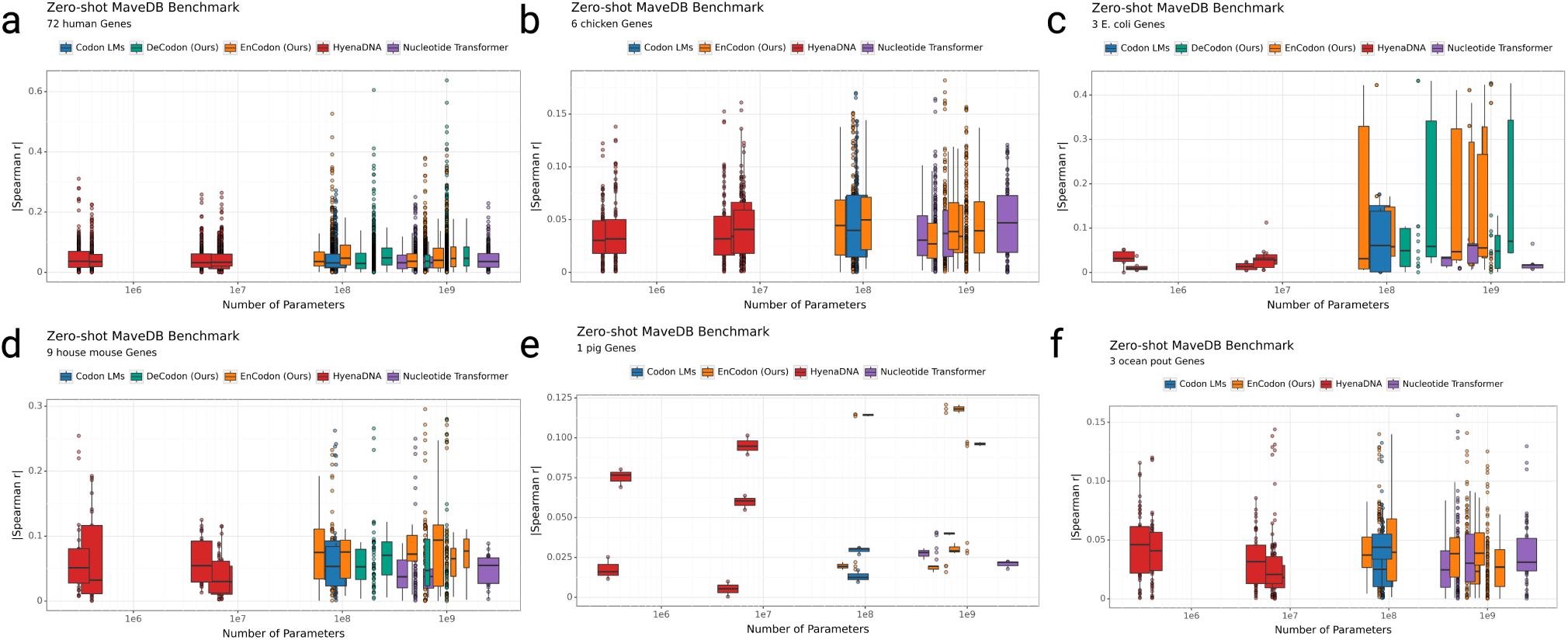
Relationship between model size (log-scaled number of trainable parameters, x-axis) and Zero-shot MaveDB performance, reported as the distribution of absolute Spearman correlations (y-axis) for each organism: **a)** Chicken, **b)** E. coli, **c)** House mouse, **d)** Pig, **e)** Ocean pout, and **f)** Fruit fly.

**Supplementary Figure 12:**
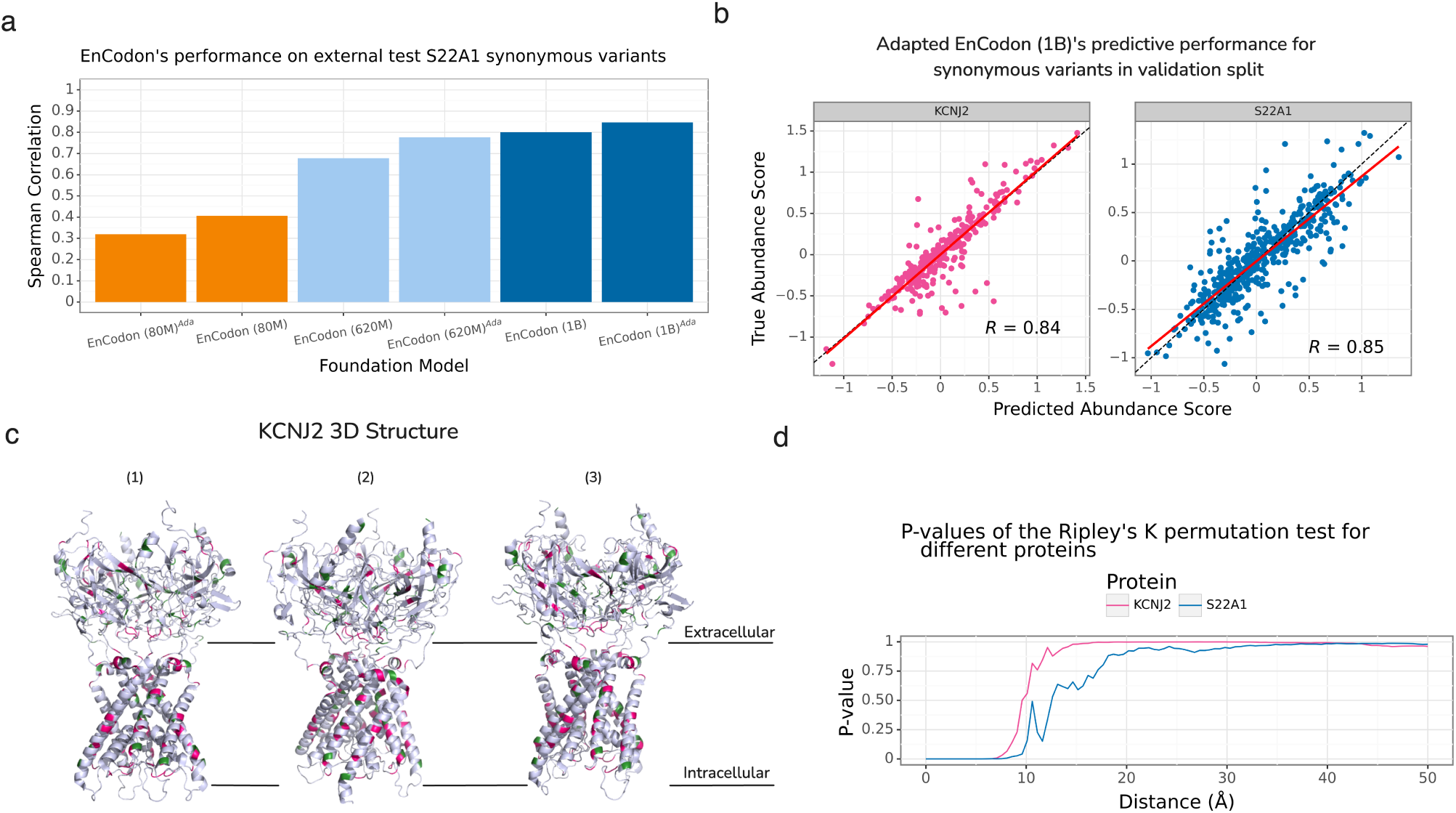
**a)** 3D structure visualization of KCNJ2 protein from 3 different angles where critical variants identified by EnCodon (1B) are colored in pink and green. **b)** Performance of EnCodon (1B)*^Ada^* in abundance prediction for KCNJ2 (left) and SLC22A1 (right) variants in the validation set. **c)** Bar plot of Speraman correlations of fine-tuned EnCodon models on the external set of SLC22A1 variants. **d)** Line plot of computed p-values at different distance cut-offs for KCNJ2 and SLC22A1 proteins.

**Supplementary Figure 13:**
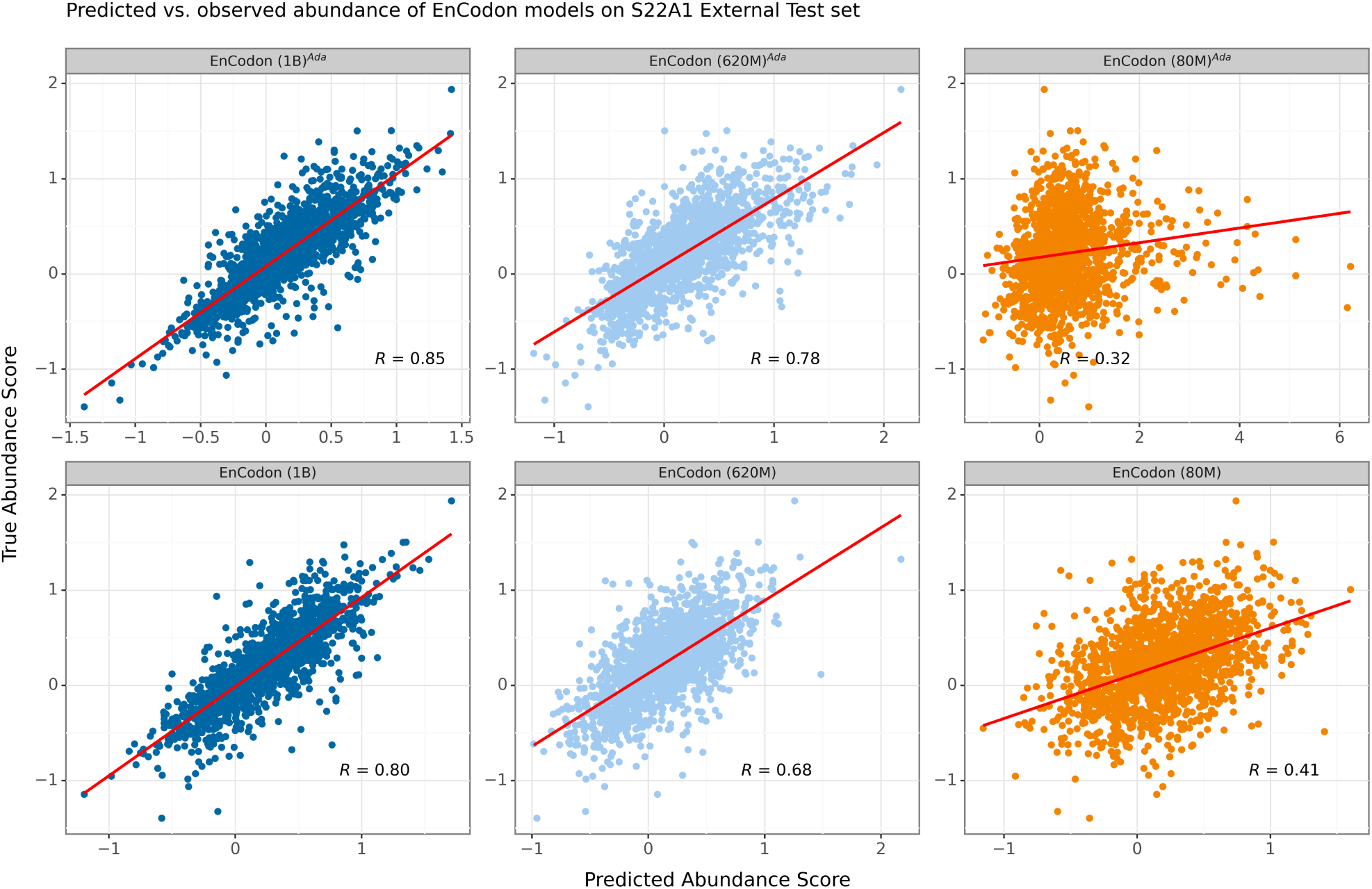
Scatter plot of all 6 fine-tuned EnCodon models on the external test set of SLC22A1 variants. 3 pre-trained (bottom row) and 3 eukaryotic adapted EnCodon models (top row) were fine-tuned.

**Supplementary Figure 14:**
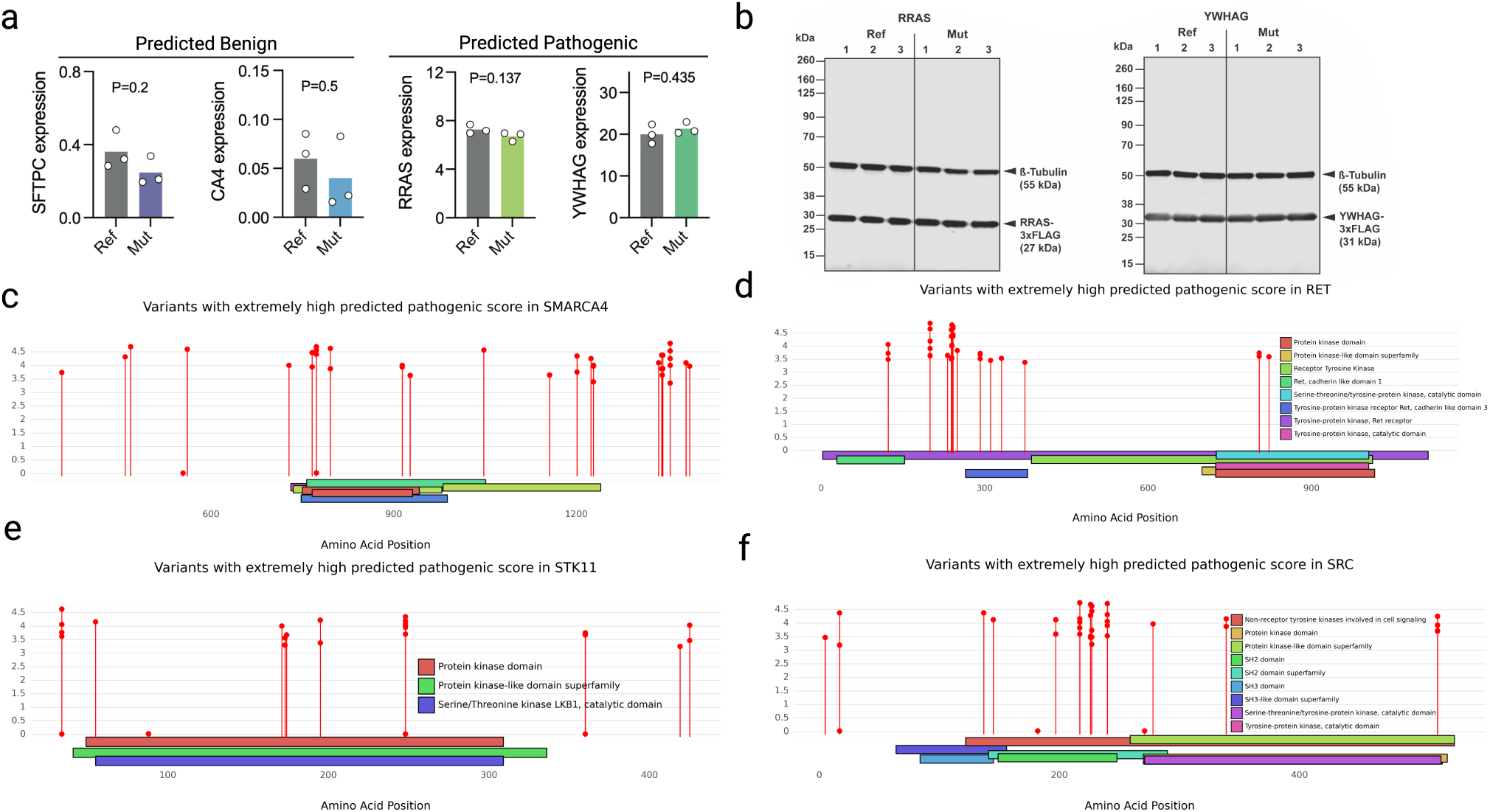
**a)** Barplot of observed gene expression levels of 4 other tested synonymous variants (2 controls and 2 predicted as pathogenic). **b)** Immunoblots of FLAG-tagged RRAS and YWHAG protein variants expressed in HEK293T cells, showing three biological replicates for both wild-type (Ref) and mutated (Mut) sequences. *β*-tubulin signal is used as a loading control. Lollipop plots showing potential synonymous variants with extremely pathogenic score for **c)** SMARCA4, **d)** RET, **e)** STK11, and **f)** SRC.

## 5 Methods

### 5.1 Pretraining data

For pretraining, we aggregated coding sequences (CDS) from all available species in the [NCBI Genomes database](https://www.ncbi.nlm.nih.gov/home/genomes/). We utilized the most recent genomic annotation files (GTF files) available for each organism to ensure accuracy and completeness. The resulting dataset, which we refer to as *NCBI CDS* data, comprises a total of 60 million sequences from over 5,000 species. This comprehensive collection serves as a robust foundation for training models on a diverse set of genetic sequences.

### 5.2 Eukaryotic adaptation data

Due to overwhelming representation of bacterial coding sequences in the NCBI CDS dataset, we curated a separate dataset of eukaryotic coding sequences for adaptation of our pre-trained models. The dataset contains 567,281 coding sequences from 227 eukaryotic species, including human, mouse, fruit fly, and zebrafish. This dataset was used to fine-tune the pre-trained models on eukaryotic sequences, enabling them to better capture the nuances of eukaryotic codon usage.

### 5.3 ClinVar dataset

We obtained the latest variant summary file in TSV format (variant_summary.txt.gz) from the ClinVar database, which initially contained 2,366,650 GRCh38-aligned variants. From this dataset, we extracted coding single-nucleotide variants (SNVs) with a review status of 2+ stars, indicating higher confidence in clinical interpretation. These SNVs were then mapped to the GRCh38 RefSeq reference genome to extract corresponding coding sequences. Two versions of the preprocessed ClinVar dataset were created: v0.1 and v0.2. Version v0.2 includes all variants from v0.1 and additional variants with uncertain clinical significance (VUS). v0.1 was employed for zero-shot benchmarking of language models, while v0.2 was used to identify candidate synonymous variants for experimental validation. Detailed statistics of the dataset versions are provided in Supplementary Table 2.

**Supplementary Table 2.**
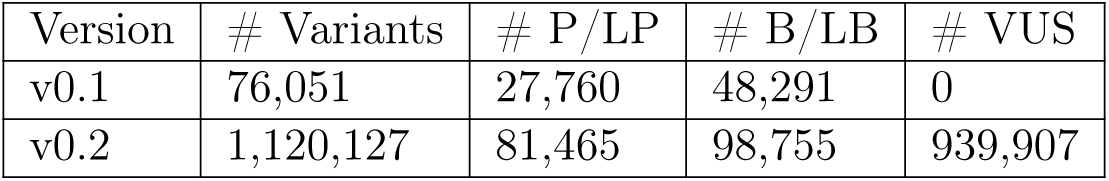
Summary statistics of the different versions of the preprocessed ClinVar dataset. P/LP: Pathogenic/Likely Pathogenic; B/LB: Benign/Likely Benign; VUS: Variant of Uncertain Significance.

### 5.4 MaveDB Collection

We retrieved 3,153 experimental datasets from the [MaveDB](https://www.mavedb.org/) database using its API, which included their corresponding score set CSV files. To focus on coding sequences, we filtered out experiments lacking nucleotide-level information for the target sequence or its variants. Additionally, we restricted the dataset to single-nucleotide variants (SNVs), resulting in a final collection of 1,911,200 variants across 13 species. The species included Human (n = 519,771), Chicken (n = 122,460), Ocean pout (n = 73,578), House mouse (n = 27,608), E. coli (n = 14,831), Thale cress (n = 14,817), Fruit fly (n = 9,192), Antarctic eel pout (n = 8,229), African clawed frog (n = 4,347), Viviparous blenny (n = 4,089), Pig (n = 3,969), Norway rat (n = 3,768), and Ocean sunfish (n = 3,639).

### 5.5 Membrane protein abundance and surface expression deep mutational scanning

Deep mutational scanning datasets for membrane proteins were generated as previously reported in more detail within manuscripts [32, 51, 89]. However, in all cases, the oligo-based library generation pipeline, DIMPLE, is used to make variant libraries, and these libraries are BxBI-mediated integrated in stable landing pad cell HEK293T cell lines [54, 55]. These library containing cell lines were into four populations by fluorescent cell sorting based on a massively parallel measure for protein abundance and/or surface expression. For the protein abundance assay, cells containing mutational libraries were sorted based on a split fluorescent protein complementation [53], whereas for surface expression screens cells were sorted based on fluorescent antibodies that recognize extracellular exposed epitopes [16]. Once these cells are separated, DNA is extracted, PCR-amplified, fragmented, and prepared for sequencing by Illumina Nextera kits and sequenced on an Illumina Novaseq 6000 short sequencer. Variant effect scores were generated using Enrich2 [69].

### 5.6 Open Reading Frame (ORF) Data

We obtained a collection of 7,264 translated open reading frames (ORFs) from the supplementary material provided by [57], which were filtered and derived from Ribo-seq studies. Specifically, we concatenated ORFs from two Excel sheets titled **S2. PHASE I Ribo-seq ORFs** and **S3. Singlestudy Ribo-seq ORFs**. The ORF locations were then mapped to the GRCh38 RefSeq genome to ensure consistency with our other datasets.

### 5.7 mRNA Stability Dataset

For the mRNA stability downstream benchmark, we utilized a preprocessed collection of mRNA decay rate datasets provided by [2]. This dataset encompasses 39 human and 27 mouse transcriptome-wide studies, collectively containing 26,725 mRNA sequences (12,981 for human and 13,744 for mouse) with reported half-life measurements. This data is crucial for understanding mRNA stability and its implications in gene expression regulation.

### 5.8 Synonymous protein variants Dataset

#### 5.8.1 Cloning of plasmids expressing synonymous protein variants

For restriction enzyme cloning, custom double-stranded DNA sequences (Twist Biosciences) encoding synonymous protein variants were designed as follows: 5’-AAGCTG-GCTAGC (NheI)-GCCACC (Kozak sequence)-Variable Coding Sequence-GGT GGA GGC GGT AGC (GGGS linker)-GACTACAAGGAC-CACGACGGCGATTATAAGGATCACGACATCGACTACAAAGACGACGATGACAAG (3xFLAG)-XXX (Gene-specific Stop Codon)-GAATTC (EcoRI)-TGCAGA-3’. The exact DNA sequences for every synonymous protein variant used for cloning can be found in Supplemntary Table (other details can be found in Supplementary Table 3). DNA was double-digested with NheI-HF (New England Biolabs, #R3131L) and EcoRI-HF (New England Biolabs, #R3101L) for 30 min at 37°C, and heat-inactivated for 20 min at 80°C. pcDNA3.1 vector (Thermo Fisher Scientific, #V79020) was double-digested with NheI-HF and EcoRI-HF for 30 min at 37°C, dephosphorylated by addition of Quick-CIP (New England Biolabs, #M0525L) (10 min at 37°C), and heat-inactivated for 20 min at 80°C. For ligation in 1x rCutSmart™ Buffer (New England Biolabs, #B6004S) supplemented with 1 mM ATP, 50 ng crude digested pcDNA3.1 vector was combined with a 7-times molar excess of crude digested insert and incubated for 30 min at room temperature after addition of 1 *µ* L Quick Ligase (New England Biolabs, #M2200). NEB stable competent E. coli (New England Biolabs, #C3040H) were transformed according to manufacturer’s instructions. Bacteria were grown on LB-agar plates supplemented with 0.1 mg/mL carbenicillin overnight at 37°C. LB supplemented with 0.1 mg/mL ampicillin was inoculated with single colonies and incubated in a shaking incubator at 37°C overnight. Plasmid DNA was extracted using ZymoPURE™ Plasmid Miniprep Kit (Zymo Research, #D4210) and eluted in nuclease-free water. The plasmid DNA concentration was measured using a NanoDrop™ One spectrophotometer (Invitrogen) by measuring the absorbance at 260 nm, and the plasmids were fully sequenced by Nanopore sequencing (Quintara Biosciences).

#### 5.8.2 Western blotting of synonymous protein variants

HEK293T cells (ATCC, #CRL-3216) were maintained in DMEM (Gibco, #11965-092) supplemented with 10% FBS (Avantor, #97068-085), under standard tissue culture conditions (37°C, 5% CO2) and were passaged every 2–3 days. To express synonymous protein variants, HEK293T cells were seeded in a 24-well format (75,000 cells per well), and after 24 hours transfected with 0.5 *µ* g variant-encoding plasmid using TransIT transfection reagent (Mirus Bio, #MIR6606). 24 hours after transfection, the cell culture media was changed, and cells were harvested after incubating for another 24 hours. Cells were lysed in RIPA buffer (Thermo Fisher Scientific, #89901) supplemented with 1x Halt protease inhibitor cocktail (Thermo Fisher Scientific, #78425) by shaking for 30 min at 4°C. After centrifugation for 30 min (4^◦^C) at 14,000 x g, the supernatant was collected and the protein concentration was determined by BCA assay (Thermo Fisher Scientific, #23225). Protein samples were stored at −20°C. After denaturing samples for 5 min at 95°C in 1x NuPAGE LDS loading buffer (Invitrogen, #NP0007) supplemented with 50 mM DTT, 5 *µ* g of total protein per sample and a pre-stained protein ladder (LI-COR Biosciences, #928-60000) were separated on a NuPage Bis-Tris Mini Protein Gel, 4–12% (Invitrogen, #NP0322BOX) for 1 hour at 120 V in 1x NuPAGE™ MES SDS running buffer (Thermo Fisher Scientific, #NP0002). Protein bands were transferred onto a PVDF membrane (Invitrogen, #IB34001) using the iBlot3 Western Blot Transfer System (Invitrogen). After blocking the membrane in SuperBlock blocking buffer (Thermo Fisher Scientific, #37537) for 10 min at room temperature, it was incubated overnight at 4°C with rabbit anti-FLAG monoclonal antibody (Cell Signaling, #14793) and mouse anti-*β*-tubulin monoclonal antibody (Sino Biological, #100109-MM05T), both diluted 1:2,000 in blocking buffer. The membrane was washed three times for 10 min each in 1x TBS (Thermo Fisher Scientific, #J62938.K3-T) supplemented with 0.1% (v/v) Tween-20 (TBS-T), followed by incubation at room temperature for 30 min with anti-rabbit IgG DyLight 800 4X PEG conjugate (Cell Signaling, #5151) and anti-mouse IgG DyLight 680 conjugate (Cell Signaling, #5470), both diluted 1:5,000 in blocking buffer. Membranes were washed two times for 5 min in TBS-T, followed by a 5 min wash in TBS without Tween-20, and the fluorescence signal was recorded on an Odyssey DLx Imager (LI-COR) and analyzed in Image Studio (LI-COR Biosciences, version 5.2.5). Protein expression levels were calculated by normalizing the anti-FLAG signal (800 nm channel) to the corresponding anti-*β*-tubulin signal (700 nm channel).

**Supplementary Table 3.** Nominated synonymous variants, including detailed PHRED scores and predictions from our codon-based model (showing pathogenicity scores and corresponding predicted labels)

| HGVS | Gene | PHRED | Prediction |
| --- | --- | --- | --- |
| NM_006412.4(AGPAT2):c.702C>T | AGPAT2 | 0.034 | -3.207 (pathogenic) |
| NM_006270.5(RRAS):c.333C>T | RRAS | 1.505 | -3.057 (pathogenic) |
| NM_002872.5(RAC2):c.501C>T | RAC2 | 0.069 | -2.928 (pathogenic) |
| NM_012479.4(YWHAG):c.564C>T | YWHAG | 0.295 | -2.667 (pathogenic) |
| NM_001317778.2(SFTPC):c.228G>C | SFTPC | 0.012 | 0.097 (benign) |
| NM_000717.5(CA4):c.492G>A | CA4 | 0.061 | 0.000 (benign) |

### 5.9 COSMIC Census Mutations Database

We acquired the Mutant Census v100 file from the [COSMIC](https://cancer.sanger.ac.uk/cosmic) database [75], which details coding mutations in genes listed in the [Cancer Gene Census](https://cancer.sanger.ac.uk/census). The raw dataset comprised 1,949,478 coding mutations. We filtered the data to retain only single-nucleotide variants (SNVs) and mapped their locations to the GRCh38 RefSeq genome. The final processed dataset included 244,400 coding variants, forming a key resource for studying mutational impacts in cancer.

### 5.10 Foundation Models

#### 5.10.1 EnCodon

##### Architecture

The architecture of EnCodon is inspired by the RoFormer as described in [76] (see 5.11). It is fundamentally based on the transformer encoder model, which incorporates multiple layers of self-attention mechanisms followed by position-wise feed-forward networks but similar to RoFormer, we use rotary positional encoding (RoPE) instead of absolute or relative PE approaches. EnCodon is a large language model designed to provide contextual codon-level hidden representations (i.e., 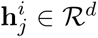 where *j* = 1*, …, L*) for any given input coding sequence (i.e., **x***^i^*) of length *L*.

The EnCodon model comprises three primary components: a learnable vocabulary embedding layer, a stack of transformer encoder blocks, and a language modeling head. The input coding sequence **x** is represented as a vector of *L* token IDs—integers that represent either a codon or a special token. The EnCodon vocabulary includes 69 tokens: 64 codons and 5 special tokens, namely [CLS], [SEP], [PAD], [MASK], and [UNK]. [CLS] and [SEP] are control tokens prepended and appended to each input coding sequence, marking the start and end of the sequence, respectively. The [PAD] token is used for padding during batched training or inference, [UNK] is used for unknown codons, and [MASK] is used for masking during training. It is important to note that the model is not provided with any information regarding the nucleotide composition of the codons; the only input information consists of token IDs. Consequently, the model does not have prior knowledge of the nucleotide differences between codons during training. Mathematically, EnCodon model takes the tokenized input sequence *x^i^*, which contains *L* token IDs:

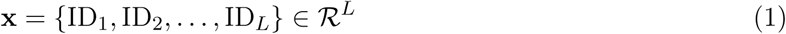

The embedding layer processes **x** to produce the embedding tensor **E**, defined as:

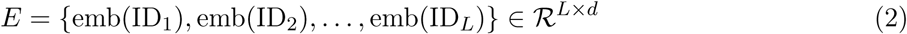

Subsequently, the embedding tensor **E** is passed through multiple transformer encoder blocks, which follow the architecture described in [76]. Each block consists of a rotary attention layer, which applies rotary positional encoding (see 5.11) followed by a self-attention operation (see 5.10.3), and a position-wise feed-forward network. We employed the “Sub-LayerNorm” approach proposed by [86] to enhance expressivity and utilized a scalable, theoretically-derived initialization strategy. Detailed hyperparameters for each pre-trained EnCodon model are provided in Supplementary Table 1.

##### Pretraining Procedure

For pretraining, EnCodon uses the Masked Language Modeling (MLM) objective, defined as:

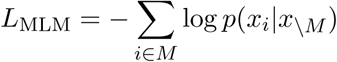

where *M* represents the set of masked positions, *x_i_*is the true token at position *i*, and *x*_\_*_M_* denotes the input sequence with tokens at positions in *M* masked. This loss function encourages the model to accurately predict the original tokens based on the surrounding context. The EnCodon foundation models were pretrained on 2 NVIDIA H100 GPUs for 2 weeks. The models were implemented using PyTorch [63] and the HuggingFace [87] framework was used for pretraining.

#### 5.10.2 DeCodon

##### Architecture

DeCodon is a controllable generative language model specifically designed for the codon-level generation of coding sequences. Similar to EnCodon, the architecture of DeCodon is similar to the RoFormer [76] but also utilizing Extrapolatable Position Embedding (XPOS) [77] in rotary attention layers (See 3a). XPOS has recently shown significant improvement in the generalization of mega-scale language models like PaLM[15] on very long sequences without sacrificing models’ performance on shorter sequences. We used sequences’ organism as the conditional information for DeCodon making the generation controllable. Similar to EnCodon, DeCodon operates on input sequences of codons, but it is optimized for sequence generation tasks rather than contextual encoding.

The DeCodon model consists of three main components: a learnable vocabulary embedding layer, a stack of transformer decoder blocks, and a sequence generation head. The input to DeCodon is a sequence of *L* token IDs, where each ID corresponds to a codon, specie, or a special token from a vocabulary that includes the 5124 tokens: 64 codons, 5 special tokens, and 5055 organism tokens. The model is autoregressive, meaning that it generates each token in the sequence one at a time, conditioned on the previously generated tokens in addition to the organism of interest.

Given an input sequence *x^i^* of length *L*:

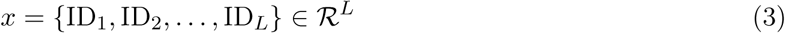

The embedding layer converts this input sequence into an embedding tensor **E**, defined as:

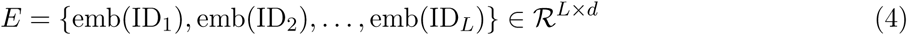

This embedding tensor is then passed through a series of transformer decoder blocks. Each block consists of a masked self-attention layer, where attention is only computed over previous tokens in the sequence, followed by a position-wise feed-forward network. The use of rotary positional encoding (see 5.11) allows the model to effectively capture the sequential dependencies between codons. As in EnCodon, we employed the “Sub-LayerNorm” approach [86] for improved expressivity and model stability. The final output is produced by the sequence generation head, which predicts the next token in the sequence based on the transformer outputs.

##### Training Procedure

DeCodon model is trained using a standard causal (i.e. autoregressive) language modeling (CLM) objective, which aims to maximize the likelihood of the target sequence given the input sequence:

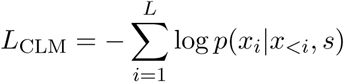

where *x_<i_* denotes the sequence of tokens preceding the *i*-th token. *s* also denotes the sequence’s specie. This loss encourages the model to generate codon sequences that are contextually and sequentially accurate. The DeCodon model was pretrained on the same dataset and hardware as EnCodon, utilizing PyTorch [63] and the HuggingFace [87] framework.

#### 5.10.3 Self-Attention Operation

The self-attention mechanism, proposed in [85], is a data-driven operation that quantifies the information flow between all possible pairs of tokens in an input sequence. This mechanism forms as the core component of the Transformer architecture.

Mathematically, let *X* = {*x*_1_*, x*_2_*, …, x_N_* } ∈ ℝ*^N^*^×^*^d^* be the input sequence of *N* token representations, where each *x_i_* ∈ ℝ*^d^*. The self-attention operator first transforms the input token representations into three different representations called query, key, and value:

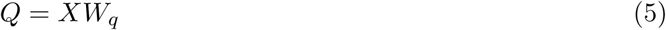

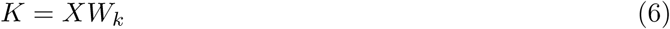

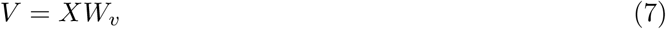

where *W_q_, W_k_, W_v_* ∈ ℝ*^d^*^×*d*^ are learned parameter matrices.

The attention scores are then computed as:

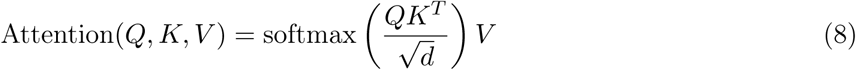

where *Q, K, V* ∈ ℝ*^N^*^×^*^d^* represent the query, key, and value matrices derived from the input representations. The scaling factor 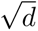 is introduced to mitigate the effect of large dot products in high-dimensional spaces.

This mechanism allows the model to weigh the importance of different tokens within the input sequence, regardless of their positions. The output of the self-attention operation is a weighted sum of the value vectors, where the weights are determined by the compatibility between the query and key vectors.

In practice, multi-head attention is often employed, which involves applying multiple sets of query, key, and value projections in parallel:

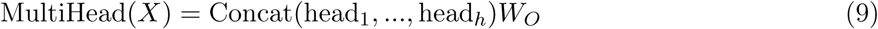

where each head is computed as:

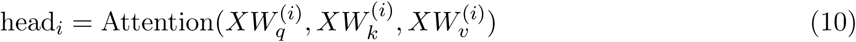

Here, 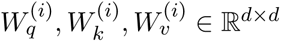 and 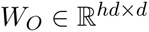 are learned parameters, and *h* is the number of attention heads.

The self-attention mechanism enables the model to capture complex dependencies and relationships within the input sequence, contributing significantly to the success of Transformer-based models in various natural language processing tasks.

### 5.11 Rotary Positional Encoding (RoPE)

Self-attention is a position-invariant operation, suggesting the need to encode positional information explicitly. Several approaches have been proposed to incorporate positional information in self-attention which can be categorized into absolute, relative, and hybrid PE methods. Rotary Positional Encodings (RoPE), recently proposed by [76], is a positional encoding (PE) approach that is a combination of relative and absolute PE approaches. RoPE has showed consistent improvement in various natural language understanding and computer vision tasks making it a common practical choice for positional information encoding which motivated us to use them in EnCodon and DeCodon architectures.

### 5.12 Baselines

#### 5.12.1 NucCNN

NucCNN is a convolutional neural network (CNN) model with a nucleotide vocabulary embeddings. We used 5 convolutional layers with kernel sizes of 3, 5, 7, 9, and 11, each followed by a max-pooling layer. The model was trained using the same dataset as EnCodon and DeCodon in the benchmarked tasks, with the same hyperparameters. The model was implemented using PyTorch and the HuggingFace framework.

#### 5.12.2 CodonCNN

CodonCNN is almost identical to NucCNN, but it uses a codon vocabulary instead of nucleotides. The model was trained using the same dataset as EnCodon and DeCodon in the benchmarked tasks, with the same hyperparameters. The model was implemented using PyTorch and the HuggingFace framework.

### 5.13 Sequence Embeddings

The proposed EnCodon and DeCodon models generate codon-level representations, where each codon in the input sequence is represented by a d-dimensional vector. For ‘sequence embedding,’ we calculate the average of all codon-level embeddings in the EnCodon models, while for the DeCodon models, we use the representation of the last codon in the sequence.

### 5.14 Fine-tuning scheme for synonymous variant prediction

To fine-tune the pre-trained EnCodon language models for the synonymous variant prediction task, we reused the pre-trained language model head without adding any new layers or parameters. Specifically, for each synonymous variant, EnCodon takes the wild-type coding sequence as input and computes the log-likelihood ratio between the wild-type and mutated codon at the variant position (See Figure 5a). This log-likelihood ratio is used as the predicted score for the variant. The model was fine-tuned using the Huber loss function, with a learning rate of 1e-5, over 5 epochs.

### 5.15 Metrics

#### 5.15.1 KNN Purity

KNN purity evaluates clustering quality by measuring the homogeneity of KNN clusters with respect to their ground-truth labels. It uses the K-nearest neighbors algorithm to assess how well cluster points match their true labels. The KNN purity score is mathematically defined as *P* :

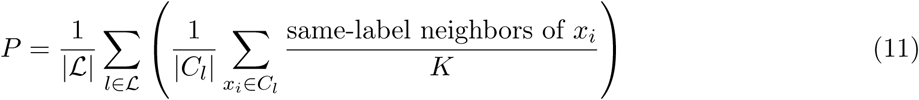

where 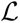 is the set of unique labels, *C_l_*is the set of points with label *l*, and *K* is the number of nearest neighbors. The score ranges from 0 to 1; a higher score indicates more homogeneity. A score of 1 means perfect clustering (all points in a cluster share the same label), while a lower score indicates greater heterogeneity.

#### 5.15.2 Spearman correlation

Spearman’s rank correlation coefficient is a non-parametric measure that assesses the strength and direction of a monotonic relationship between two variables. Mathematically, for two variables X and Y, the Spearman correlation coefficient, *ρ*, is given by the formula:

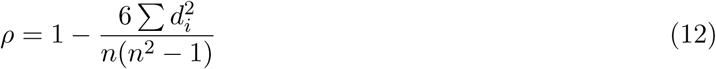

where *d_i_* = *rank*(*X_i_*) *− rank*(*Y_i_*) is the difference between the ranks of each pairs of values and n is the number of data points. It is important to mention that this correlation is robust to outliers and skewed data, making it suitable for datasets where traditional parametric assumptions do not hold.

#### 5.15.3 Pearson correlation

Pearson correlation coefficient, *r*, is a measure of linear correlation between two random variants *X* and *Y* :

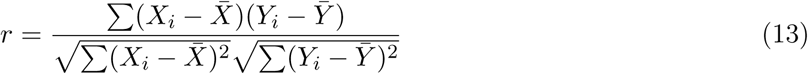

where *X_i_*, and *Y_i_* are individual data points and *X̅* and *Y̅* are sample averages of samples drawn from *X* and *Y*, respectively. The correlation ranges from −1 (negative linear relationship) to +1 (positive linear relationship) and 0 indicates no linear relationship.

#### 5.15.4 Synonymous Codon Confusion

Synonymous Codon Confusion is a amino acid level metric, measuring the how often a langauge model incorrectly choose synonymous codon over each other. The metric can be computed directly from the confusion matrix of a codon-level language model. Mathematically speaking:

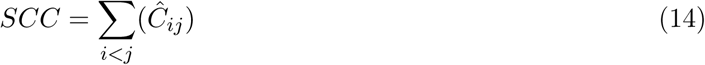

where *Ĉ* is the row-normalized confusion matrix of a codon-level lagnuage model.

#### 5.15.5 variant effect predicted score (VEP)

Focusing on the language models zero-shot applicability on variant effect prediction tasks, we used different formulations for encoder and decoder langauge models. Specifically, concerning encoder models, we computed the masked lanuage modeling (MLM) pseudolikelihood ratio between wild-type and mutated codon/nucleotide/token at the position of mutation. Mathematically speaking, we formulate VEP as follows:

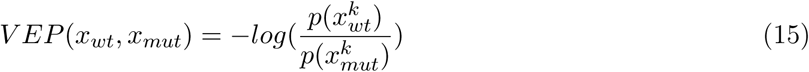

where *x_wt_* and *x_mut_* are wild-type and mutated coding sequences, respectively. k indicates the variant’s token-level position in the coding sequence depending on the tokenization used for the encoder language model. For example, we used variant’s codon position to compute the log-likelihood ratio (i.e. VEP) score with our EnCodons.

Concerning decoder language models, we reported the difference between sequence likelihood scores (*SL*) computed for the wild-type and mutated coding sequences. In constrast to encoder language models, we don’t take the variant’s location information into account for computing the variant effect predicted (VEP) score:

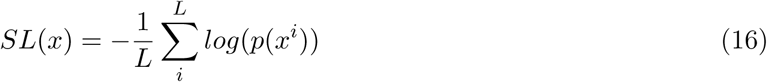

where *x^i^* denotes the i-th position in the input sequence of length L and *p*(.) is the output probability at the i-th position computed by the decoder lanaguge model. Finally, we reported the difference between the two computed sequence likelihoods (*SL*) as the predicted score:

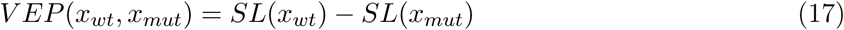

### 5.16 Spatial Clustering Analysis Using Ripley’s *K* Functions (*L*)

To investigate the spatial distribution of extreme variants within the 3D structure of various proteins, we applied Ripley’s *K* function, a statistical method designed to detect clustering or dispersion of points in a spatial domain [67]. Ripley’s *K* function in three dimensions is defined as:

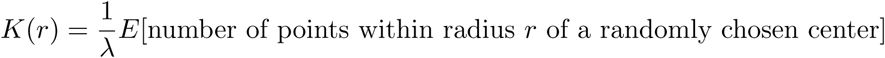

where *λ* represents the density of points, *r* is the distance threshold, and *E*[*·*] denotes the expectation. For each protein in the experiment, we performed a radius range *r* from 0 to 50 Å with 100 discrete intervals. The statistical significance of clustering was assessed using a permutation test with 10,000 iterations. Specifically, we generated 10,000 sets of randomly chosen variants (*V_r_*) as the “null distribution” with the same number of points as the observed “critical” synonymous variants (*V_c_*) for the protein. Then, we first extracted the C*α* atom coordinates of the amino acids of each set of points from the protein PDB structure using the Bio.PDB module in Python. Next, we computed pairwise Euclidean distances between the C*α* atoms and calculated normalized Ripley’s *K* function at various radius *r* using:

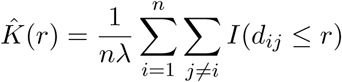

where *n* is the total number of points, and *I*(*·*) is an indicator function that equals 1 if the condition inside is true. We calculated Ripley’s *K* function at 10000 different distances ranging from 0 to 100Å. Finally, The observed *K* values were compared against the null distribution to compute p-values, representing the probability that the observed clustering occurred by chance:

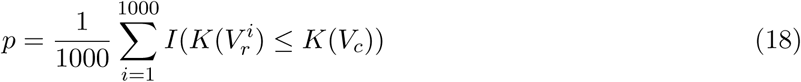

